# A thalamostriatal brake counteracts cortical recruitment of striatal ensembles in levodopa-induced dyskinesia

**DOI:** 10.64898/2026.07.22.740223

**Authors:** Jie Wang, Xin-Yu Tu, Jing-Xue Liang, Jiu-Yang Sun, Wenying Xu, Yong-Ling Wang, Xin Yi, Yan-Jiao Wu, Xin-Ni Li, Ying-Xiao Liu, Zhenguo Liu, Wei-Guang Li, Lu Song

## Abstract

Levodopa-induced dyskinesia (LID) is a disabling complication of Parkinson’s disease therapy, yet how upstream circuits recruit and restrain dyskinesia-linked striatal ensembles remains unclear. Using FosTRAP-based ensemble access in a unilateral 6-hydroxydopamine mouse model, we identified secondary motor cortex (M2) and parafascicular thalamus (PF) as dominant afferents with opposing functions. Projection-wide M2 activation promoted dyskinesia, whereas PF activation suppressed ongoing dyskinesia and shifted behavior toward non-dyskinetic states. Chronic levodopa reduced overall presynaptic terminal abundance while preserving putative contacts onto ensemble neurons, thereby increasing effective pathway-to-ensemble coupling. This remodeling followed distinct pathway rules: M2 contacts became spatially dispersed and biased toward NMDAR-mediated excitation, whereas PF inputs recruited stronger polysynaptic inhibition. Dyskinesia preferentially re-engaged ensemble-projecting M2 neurons, but reactivated PF neurons were topographically segregated from PF neurons directly innervating the ensemble. Accordingly, selective M2-to-ensemble stimulation promoted dyskinesia, whereas selective PF-to-ensemble stimulation was ineffective. Finally, ensemble-restricted *Grin1* knockdown reduced peak dyskinesia and weakened M2-driven dyskinesia. These findings define LID as a targetable imbalance between cortical ensemble recruitment and thalamostriatal restraint.

## Introduction

Parkinson’s disease (PD) is a progressive neurodegenerative disorder caused by degeneration of midbrain dopamine neurons and characterized by bradykinesia, rigidity, tremor, and postural instability.^1–3^ Levodopa remains the most effective pharmacological therapy for restoring movement, yet long-term treatment is frequently limited by levodopa-induced dyskinesia (LID), a disabling syndrome of abnormal involuntary movements that can become as burdensome as parkinsonism itself.^4–8^ In some cohorts, dyskinesia develops in the majority of levodopa-treated patients, reaching ∼80% within five years.^2^ This complication progressively narrows the therapeutic window: doses required to relieve parkinsonian motor impairment increasingly provoke dyskinesia, whereas dose reduction often worsens mobility and quality of life. Although dose fractionation and adjunct medications can provide symptomatic benefit in some patients, mechanism-based strategies that suppress dyskinesia while preserving levodopa efficacy remain limited.

At the circuit level, LID is widely viewed as a maladaptive plasticity disorder within basal ganglia loops, with the dorsal striatum serving as a key substrate where dopamine denervation and pulsatile dopamine replacement converge. Dopamine depletion alters receptor coupling, intracellular signaling, and plasticity thresholds, whereas repeated intermittent levodopa exposure unmasks and amplifies these latent adaptations.^9–13^ A large body of work has linked dyskinesia to biased recruitment of direct-over indirect-pathway striatal projection neurons, together with abnormal glutamatergic transmission and plasticity at corticostriatal and thalamostriatal synapses.^14–16^ Yet an essential translational question remains unresolved: if dopamine replacement is diffuse and excitatory afferents to the striatum are dense and heterogeneous, which upstream pathways repeatedly push striatal networks into a dyskinetic state, and which pathways can restrain this pathological recruitment?

Recent work has sharpened this question by reframing LID as an ensemble-based phenomenon rather than a uniform property of all direct-pathway medium spiny neurons.^17,18^ Activity-dependent genetic access using FosTRAP identifies a stable, non-random subpopulation of striatal neurons that is repeatedly recruited during dyskinesia and enriched for direct-pathway neurons; reactivating these tagged neurons is sufficient to evoke dyskinesia even without levodopa, whereas inhibiting them attenuates LID.^17^ Complementary *in vivo* recordings and imaging similarly reveal sparse striatal neuronal populations with exaggerated levodopa-evoked activity that track dyskinesia severity and encode distinct dyskinetic motifs.^16,19^ These studies identify the downstream striatal population that contributes to dyskinesia, but they leave open the upstream logic of its recruitment. In particular, the identity of the afferent pathways that access dyskinesia-linked ensemble neurons, how chronic levodopa remodels those inputs, and whether different upstream circuits drive or restrain dyskinesia remain poorly understood.^20^

This gap is especially important for circuit-targeted therapeutic strategies, because anatomical connectivity does not necessarily identify the circuit module that controls a pathological state. The cortico–basal ganglia–thalamic network is organized into parallel, topographically matched subnetworks,^21,22^ raising the possibility that dyskinesia-related activity might remain constrained within anatomically defined channels. Here, we tested this assumption by combining activity tagging of dyskinesia-linked striatal ensembles with monosynaptic input mapping, synapse-resolved anatomy, *ex vivo* electrophysiology, 3D behavioral analysis, and causal circuit perturbations. We identify secondary motor cortex (M2) and the parafascicular nucleus of the thalamus (PF) as dominant afferents to dyskinesia-linked ensemble neurons with opposing effects on behavior: M2 activation exacerbates dyskinesia, whereas PF activation suppresses ongoing dyskinesia and shifts behavior toward non-dyskinetic states. Chronic levodopa increases effective pathway-to-ensemble coupling despite terminal loss, but imposing distinct synaptic mechanisms—NMDAR-weighted excitation for M2 inputs and stronger polysynaptic inhibition for PF inputs. Finally, we show that dyskinesia preferentially engages ensemble-projecting M2 neurons, whereas PF displays a topographic dissociation between dyskinesia-linked activity and direct ensemble connectivity, explaining why broad PF stimulation is anti-dyskinetic whereas selective PF-to-ensemble activation is ineffective. Together, these findings identify LID as a targetable circuit imbalance between an NMDAR-weighted motor cortical drive and an ensemble-external parafascicular thalamostriatal brake.

## Methods

### Study design

This study was designed to identify upstream circuit mechanisms that recruit or suppress dyskinesia-linked striatal ensembles in a mouse model of LID. Adult male mice were used to establish unilateral dopamine depletion, induce dyskinesia by repeated levodopa administration, tag dyskinesia-engaged striatal neurons using FosTRAP, map their upstream inputs, and test the behavioral and synaptic consequences of manipulating cortical and thalamostriatal pathways. Experimental approaches included stereotaxic viral delivery, unilateral 6-hydroxydopamine (6-OHDA) lesioning, activity-dependent genetic tagging, monosynaptic rabies tracing, optogenetic and chemogenetic manipulation, fiber photometry, *ex vivo* whole-cell electrophysiology, immunohistochemistry, synapse imaging, and 3D behavioral analysis.

Animals were assigned to experimental groups in a balanced manner across batches whenever feasible. Investigators were blinded to group identity during dyskinesia scoring, offline behavioral analysis, and image quantification whenever compatible with the experimental design. Pre-established exclusion criteria are described below.

### Animals

C57BL/6J mice and transgenic reporter/driver lines on a C57BL/6J background were used. C57BL/6J wild-type mice were purchased from GemPharmatech (Nanjing, China; RRID: IMSR_JAX:000664). Fos^2A-iCreERT2^ mice (TRAP2; The Jackson Laboratory; STOCK *Fos^tm2.1(icre/ERT2)Luo^/*J; stock #030323; RRID: IMSR_JAX:030323), Ai14 reporter mice (*B6.Cg-Gt(ROSA)26Sor^tm14(CAG-tdTomato)Hze^*/J; The Jackson Laboratory; stock #007914; RRID: IMSR_JAX:007914), Ai162 reporter mice (*B6.Cg-Igs7^tm162.1(tetO-GCaMP6s,CAG-tTA2)Hze^*/J; The Jackson Laboratory; stock #031562; RRID: IMSR_JAX:031562), and LSL-H2B-GFP reporter mice (*B6.Cg-Gt(ROSA)26Sor^tm8(CAG-HIST1H2BB/EGFP)Zjh^*/J; The Jackson Laboratory; stock #036761; RRID: IMSR_JAX:036761) were used.

Double-heterozygous TRAP2::Ai14, TRAP2::Ai162, and TRAP2::H2B-GFP mice were generated by crossing TRAP2 mice with Ai14, Ai162, or LSL-H2B-GFP mice, respectively. TRAP2::Ai14 mice were used for activity tagging/reactivation analyses, *ex vivo* electrophysiology, and synapse imaging; TRAP2::Ai162 mice were used for fiber photometry; and TRAP2::H2B-GFP mice were used for rabies-based input mapping. Wild-type and/or TRAP2 mice were used for projection-wide optogenetic/chemogenetic manipulation and associated tracing experiments as indicated in the corresponding figures.

Mice were group-housed, 2–5 animals per cage, under a 12 h light/dark cycle with food and water available *ad libitum*. Unless otherwise specified, experiments were performed in adult male mice aged 3–6 months. Mice used for *ex vivo* electrophysiology were 4–8 months old.

### Virus preparation

Recombinant AAV vectors and rabies viruses were obtained from commercial sources. Most AAV vectors used in this study were purchased from BrainVTA (China), including Cre-dependent optogenetic inhibition constructs (EF1α-DIO-eNpHR3.0-EYFP) with matched fluorophore controls (EF1α-DIO-EYFP), optogenetic activation vectors for projection-wide stimulation [hSyn-driven ChR2(E123T/T159C)-mCherry with matched controls], Chronos-EGFP vectors for *ex vivo* input isolation, chemogenetic vectors (EF1α-DIO-hM3Dq-mCherry and DIO-mCherry control), retrograde Cre delivery vectors (rAAV-retro-hSyn-Cre-EGFP), and rabies helper AAVs (EF1α-DIO-H2B-EGFP-T2A-TVA and EF1α-DIO-oRVG). For synapse imaging, presynaptic bouton reporters (hSyn-Synaptophysin-EGFP and hSyn-Synaptophysin-mTagBFP2) were used; Synaptophysin-mTagBFP2 was produced by BrainVTA. EnvA-pseudotyped CVS-N2c ΔG rabies viruses (RV-CVS-EnvA-N2c(ΔG)-tdTomato for monosynaptic tracing and RV-CVS-EnvA-N2c(ΔG)-ChR2-dsRed for rabies-dependent circuit-restricted optogenetics) were also obtained from BrainVTA. For ensemble-restricted NMDAR knockdown, the shRNA sequences targeting mouse *Grin1* (5’-GGTGCAAGTG GGCATCTACAA-3’) and scrambled sequence were cloned into vector AAV2/9-EF1α-DIO-(EGFP-U6)shRNA-WPRE-hGH polyA by Shanghai SunBio Biomedical Technology (China). Vendor-reported titers were ≥ 2 × 10^12^ vg/mL for most AAVs, ≥ 5 × 10^12^ vg/mL for synaptophysin reporters, and 2 × 10^8^ IFU/mL for rabies viruses. Viral aliquots were stored at −80 °C, thawed on ice immediately before use, and not refrozen.

### Stereotaxic surgery, 6-OHDA lesioning, and viral delivery

All stereotaxic surgeries were performed under general anesthesia induced by intraperitoneal Zoletil50 (50 mg/kg) and xylazine (5 mg/kg). Animals were fixed in a stereotaxic apparatus (model 68804, RWD, China) and maintained on a thermostatically controlled heating pad. Ophthalmic erythromycin ointment was applied to prevent corneal drying. The scalp was incised, the skull was leveled, and burr holes were drilled above the target coordinates.

To generate unilateral dopamine depletion, 6-hydroxydopamine (6-OHDA; Cat#H4381, Sigma-Aldrich, USA) was prepared fresh on the day of surgery because of rapid oxidation. 6-OHDA was dissolved in ice-cold sterile 0.9% saline containing 0.2% (w/v) ascorbic acid (Cat#A4544, Sigma-Aldrich, USA) to a final concentration of 3.5 mg/ml, protected from light (foil-wrapped), and kept on ice throughout use. Mice received 1.0 μl 6-OHDA into the medial forebrain bundle (MFB) at 300 nl/min; control mice received the same volume of vehicle. For viral delivery, AAV vectors were injected via pulled glass micropipettes at 100 nl/min. In most experiments, 6-OHDA lesioning and viral injections were performed in the same surgical session. After each infusion, the pipette was left in place for 10–15 min and withdrawn slowly to minimize reflux.

Incisions were rinsed with sterile saline and sutured. Postoperative supportive care included warmed recovery cages, subcutaneous sterile saline (1 ml as needed), and softened food/nutritional supplements for up to 2 weeks. Unless specified, subsequent experiments were initiated ≥ 3 weeks after lesion/virus surgery.

Stereotaxic coordinates (mm relative to Bregma; DV relative to dura): MFB: AP − 0.7, ML −1.2, DV −4.7; M2^1^: AP +2.2, ML −1.0, DV −1.8 (300 nl); M2^2^: AP +1.4, ML −0.9, DV −1.6 (200 nl); PF: AP −2.1, ML −0.8, DV −3.35 (200 nl); CPu: AP +0.8, ML −2.2, DV −3.3 (300 nl); CPu^1^: AP +1.0, ML−2.2, DV −3.3 (600 nl); CPu^2^: AP 0.0, ML −2.7, DV −3.3 (600 nl).

### Optical fiber implantation and verification

For *in vivo* optogenetic experiments, optical fibers (200 μm core, 0.37 NA; Newdoon, China) were implanted 3–5 weeks after the initial surgery unless otherwise specified. The scalp was reopened and fibers were positioned above target coordinates (M2^1^: AP +2.2, ML −0.7, DV −1.3, leaned 10° to left; M2^2^: AP +1.0, ML −0.9, DV −1.2, leaned 10° to right; PF: AP −2.1, ML −0.8, DV −3.0; CPu: AP +0.8, ML −2.2, DV −2.7; CPu^1^: AP +1.0, ML −2.2, DV −2.7, leaned 10° to left; CPu^2^: AP 0.0, ML −2.9, DV−2.9). Three skull screws were anchored for stability and implants were secured with dental acrylic. Mice recovered for ≥ 7 days before testing.

All injection sites, viral expression patterns, and fiber placements were verified histologically. Animals were excluded if viral expression was absent/off-target or if fiber tracks were outside the intended boundaries.

### Pharmacology and activity-dependent TRAP induction

Levodopa (Cat#D1507, Sigma-Aldrich, USA) and benserazide (Cat#B7283, Sigma-Aldrich, USA) were dissolved in 0.9% sterile saline containing 0.2% ascorbic acid, protected from light, and used within 40 min of preparation. Mice received levodopa and benserazide at 6 mg/kg and 12 mg/kg, respectively, by subcutaneous injection. The low-dose levodopa solution was obtained by only adjusting the levodopa dosage to 2mg/kg. Levodopa priming began 3–4 weeks after 6-OHDA lesioning and continued 5–6 days per week.

For activity-dependent TRAP induction, mice were screened by AIM scoring after 2 weeks of levodopa priming to exclude animals with insufficient dyskinesia (see AIM inclusion criteria below). 4-hydroxytamoxifen (4-OHT; Cat#H6278, Sigma-Aldrich, USA) was administered intraperitoneally at 50 mg/kg 40 min after levodopa injection to capture dyskinesia-activated neurons. 4-OHT was first dissolved in 100% ethanol (20 mg/ml; vortexed ∼30 min), mixed with corn oil (Cat#C8267, Sigma-Aldrich, USA) and thoroughly vortexed, and ethanol was evaporated using a centrifugal concentrator (Eppendorf, Germany) for 30 min to yield a 10 mg/ml working solution in oil.

For chemogenetic experiments, clozapine-*N*-oxide (CNO; Cat#HY-17366, MedChemExpress, USA) was dissolved in sterile 0.9% saline (1 mg/ml stock, stored at −20 °C), thawed and diluted on the day of use, and injected intraperitoneally at 5 mg/kg 40 min before open-field testing or before levodopa injection.

### Behavioral testing and analysis

Behavioral experiments were performed during the light phase (08:00–20:00) in a temperature- and noise-controlled room. Mice were acclimated for ≥ 30 min before testing. Apparatus surfaces were cleaned with 75% ethanol and air-dried before the first run and between animals. Levodopa experiments were conducted at most once per day per animal; when multiple assays were performed on the same day, open field testing was performed ≥ 2 h before other procedures.

Open field activity was assessed in a 40 × 40 × 40 cm arena. Mice were placed in the center and tracked using EthoVision XT 15.0 (Noldus, Netherlands). Recording duration depended on the experiment (e.g., 10 min for parkinsonian assessment; 20 min for chemogenetic testing and NMDAR knockdown assessment; 21 min for optogenetic testing). Rotarod performance was tested on an accelerating rod (model 47650, Ugo Basile, Italy) from 4 to 40 rpm over 4 min. Passive rotation for > 1 full turn was scored as a fall. Each mouse completed three trials per day (15 min inter-trial interval) for 3 consecutive days; performance was quantified as the mean latency to fall on day 3 (days 1–2 served as training).

LID was quantified using an established abnormal involuntary movement (AIM) framework that scores axial, limb, and orolingual dyskinesia on both temporal and amplitude scales.^23,24^ Mice were habituated for 15 min in a transparent acrylic chamber (20 × 20 cm) with a rear mirror for full-body observation and video recorded for offline scoring. Levodopa was administered and AIMs were scored at 20-min intervals until dyskinesia resolved. At each time point, animals were observed for 1 min and axial, limb, and orolingual components were scored. For each subtype, a basic severity score (0–4) and an amplitude score (0–4) were assigned; the product of these scores was summed to yield a global AIM score (maximum 48). Mice were included for ensemble experiments if, after 2 weeks of priming, they reached a basic severity score ≥ 2 in at least two AIM subtypes in at least two monitoring epochs per session.

For 3D motion capture and unsupervised kinematic clustering, behavior was recorded in the BehaviorAtlas 3D-AI system (BayONE Scientific, China) using four calibrated cameras.^25^ Mice were recorded individually in a transparent cylinder (50 cm diameter, 40 cm height) and allowed to explore freely. Videos were processed in BehaviorAnalyzer for 3D reconstruction and clustering, and Behaviorexplorer was used for manual annotation and consolidation as described in the Results.

### Fiber photometry acquisition and analysis

Fiber photometry was performed ≥ 1 week after fiber implantation. Implanted ferrules were connected to a fiber photometry system (ThinkerTech, China) using a 200 μm, 0.37 NA patch cord (Newdoon, China). Excitation was time-division multiplexed at 405 nm (reference/isosbestic), 470 nm (GCaMP), and 555 nm (tdTomato). Signals were acquired at 120 Hz total (40 Hz per channel) with synchronized video recording.

Before recording, mice were acclimated for ≥ 30 min. Patch cords were pre-illuminated with high-intensity blue light for 30 min to reduce autofluorescence. Power at the fiber tip was adjusted to ∼5 μW (405 nm), ∼20 μW (470 nm), and ∼20 μW (555 nm). A 1-min dark recording (patch cord wrapped in foil) was collected to estimate channel offsets. Dyskinesia onset/offset was identified from synchronized video. For onset-aligned analyses, 3 min before and 1 min after onset were analyzed; for offset-aligned analyses, 1 min before and 3 min after offset were analyzed.

Signals were processed in MATLAB (R2022b). The 405 nm trace was linearly fit to the 470 nm (or 555 nm) trace to model motion/bleaching components and subtracted to yield motion-corrected signals. Dark offsets were accounted for using the dark recording. ΔF/F was computed relative to baseline F0 (mean of the first 2 s of the analyzed segment) after offset correction. Peak analyses were performed on 1 min windows around onset/offset; peaks exceeding mean (ΔF/F) + 3 SD were counted to quantify peak frequency and amplitude.

### Optogenetic manipulation

Optogenetic stimulation was delivered via implanted fibers connected through a rotary commutator (ThinkerTech, China) to minimize cable torsion. Blue (470 nm) and yellow (589 nm) lasers were controlled by an optogenetic system (Newdoon, China), and power at the fiber tip was measured before each experiment (Sanwa LP10, Japan). For inhibition, continuous 589 nm light (15 mW) was delivered to eNpHR3.0-expressing striatal TRAPed neurons. For activation, blue pulses (5 ms, 20 Hz) were delivered with circuit-specific powers (M2→CPu: 10 mW; PF→CPu: 15 mW; rabies-defined M2/PF→CPu^TRAPed^ inputs: 6 mW unless otherwise specified). In levodopa experiments, stimulation was delivered during the peak dyskinesia window (typically 20–60 min post-injection). Sessions consisted of three OFF–ON cycles (open field: 180 s OFF/180 s ON; AIM and BehaviorAtlas: 60 s OFF/60 s ON), and outcomes were averaged across ON and OFF epochs within each session.

### *Ex vivo* electrophysiological recordings

Acute coronal striatal slices were prepared from adult mice (4–8 months). On the day of recording, mice received levodopa (6 mg/kg) and were deeply anesthetized 30–60 min later. Animals were transcardially perfused with ice-cold, carbogenated sucrose-based slicing solution and the brain was rapidly removed into the same solution.

The slicing solution contained (in mM): 213 sucrose, 2.5 KCl, 1.25 Na_2_HPO_4_, 26 NaHCO_3_, 10 D-glucose, 2 MgSO_4_, and 2 CaCl_2_ (pH 7.3 when bubbled with 95% O_2_/5% CO_2_; ∼350 mOsm). Coronal slices (300 μm) containing dorsal striatum were cut on a Leica VT1200S in ice-cold oxygenated slicing solution, incubated in oxygenated ACSF at 34 °C for 40 min, and held at room temperature for ≥ 30 min before recording. ACSF contained (in mM): 125 NaCl, 2.5 KCl, 1.25 Na_2_HPO_4_, 25 NaHCO_3_, 12.5 D-glucose, 2 CaCl_2_, and 1 MgCl_2_ (pH 7.3; ∼300 mOsm) and was continuously carbogenated. Recordings were performed in a submersion chamber superfused with carbogenated ACSF at room temperature (22–24 °C).

Patch pipettes (3.5–5 MΩ) were filled with a Cs-based internal solution (in mM): 132.5 Cs-gluconate, 17.5 CsCl, 2 MgCl_2_, 0.5 EGTA, 10 HEPES, 4 MgATP, and 5 QX-314 (pH 7.25–7.30; 270–280 mOsm). Cells were visualized under IR-DIC optics (BX51WI, Olympus, Japan). TRAPed neurons were identified by tdTomato fluorescence; non-TRAPed neurons were recorded from neighboring tdTomato-negative cells within the same dorsolateral striatal region. Data were acquired with an Axopatch 200B, filtered at 2 kHz, digitized at 20 kHz (Digidata 1550B with HumSilencer, Molecular Devices, USA), and collected using pClamp 11. Recordings were included only if access resistance remained < 30 MΩ and stable.

Spontaneous EPSCs were recorded at −70 mV and spontaneous IPSCs at 0 mV (3 min traces; analyses performed on a fixed 50–150 s window). For pathway-specific responses, Chronos-expressing terminals were stimulated using a 473 nm LED through the objective (3–5 ms pulses; ≥ 8 sweeps; ≥ 10 s inter-sweep interval). Light intensity was adjusted to yield reliable responses while maintaining comparable conditions across cells (2–18 mW/mm^2^). oEPSCs were recorded at −70 mV and oIPSCs at 0 mV; the evoked E/I ratio was calculated as peak oEPSC amplitude divided by peak oIPSC amplitude in the same cell. Paired-pulse ratio was measured with paired stimuli separated by 50 ms and calculated as the second peak amplitude divided by the first peak amplitude. This measure was used to assess short-term synaptic dynamics during repeated pathway stimulation. NMDAR contribution was quantified at +40 mV as the current amplitude 50 ms after stimulation, and NMDA/AMPA ratio was computed as *I*_NMDA_/*I*_AMPA_, where *I*_AMPA_ was the peak oEPSC at −70 mV. To test for polysynaptic inhibitory routing, TTX (1 μM; TETRODOTOXIN, China) was bath applied to block action potential–dependent transmission, followed by co-application of TTX and 4-AP (200 μM; Cat#HY-B0604, MedChemExpress, USA). oIPSC amplitudes were quantified after ≥ 10 min wash-in per condition; currents abolished by TTX and not restored by TTX plus 4-AP were interpreted as arising from multi-neuron circuitry rather than monosynaptic release.^26^

### Immunohistochemistry and imaging

Mice were deeply anesthetized and perfused with PBS followed by 4% PFA. Brains were post-fixed overnight at 4 °C, sectioned coronally at 40 μm (VT1200S, Leica, Germany), and stored at 4 °C in cryoprotectant for up to 3 months. Free-floating sections were washed in PBS and blocked/permeabilized for 2 h in 10% normal donkey serum with 0.3% Triton X-100 in PBS. Primary antibodies were applied overnight at 4 °C with gentle agitation, followed by PBST (0.1% Tween-20) washes and 2 h incubation with Alexa Fluor–conjugated secondary antibodies at room temperature. DAPI (Cat#10236276001, Sigma-Aldrich, USA) counterstaining (10 min) was included for general histology and omitted for PSD-95 synapse imaging to avoid spectral interference. For routine verification of viral expression and fiber placement, DAPI-only staining was used.

Primary antibodies included rabbit anti-TH (1:200; RRID: AB_297840; Cat#AF-2185, Beyotime, China), rabbit anti-c-Fos (1:500; RRID: AB_2247211; Cat#2250S, Cell Signaling Technology, USA), and rabbit anti-PSD-95 (1:1,000; RRID: AB_3076111; Cat#3450S, Cell Signaling Technology, USA). Secondary antibodies included donkey anti-rabbit Alexa Fluor 488 (1:500; RRID: AB_2819209; Cat#4412S, Cell Signaling Technology, USA) and 647 (1:500; RRID: AB_3096971; Cat#4414S, Cell Signaling Technology, USA).

Widefield images were acquired on Olympus VS120/VS200 slide scanners (10× or 20×). Sections were aligned to the Paxinos and Franklin mouse brain atlas (4^th^ edition, 2013) in Fiji/ImageJ. Functional striatal domains were defined using a published parcellation map.^21^ For each region of interest, the following measures were quantified in ImageJ using consistent thresholds and segmentation parameters within an experiment: (i) area, (ii) DAPI^+^ nuclei counts (to estimate total cell number), (iii) counts of marker-positive cells (e.g., c-Fos^+^, tdTomato^+^, RV reporter^+^), and (iv) mean fluorescence intensity and intensity profile when appropriate. Co-localization was defined at the single-cell level by overlap of nuclear/cellular signals within the same soma (validated by manual inspection in representative samples). Cell densities and proportions were computed per section and then averaged across sections for each animal; regional input fractions were computed by summing counts across all analyzed sections for each region and dividing by the total number of labeled input cells across the brain.

For whole-brain rabies input mapping, one-in-six coronal sections (40 μm thickness; 240 μm spacing) spanning Bregma +2.65 mm to −5.07 mm were processed and quantified. For dorsal striatum-focused analyses, one-in-six sections spanning Bregma +1.69 mm to −1.67 mm were used. For quantification of TRAPed neuron density across striatal domains, three representative rostrocaudal levels (Bregma +1.33, +0.13, and −0.59 mm) were analyzed per mouse. For fluorescence intensity distribution analyses in M2 and PF, representative sections at Bregma +1.69 mm (M2) and −2.27 mm (PF) were used.

To quantify reactivation of striatal TRAPed ensemble neurons across dyskinesia episodes, we computed: TRAPed neuron reactivation rate = *N* (tdTomato^+^c-Fos^+^) / *N* (tdTomato^+^); Chance overlap (independence assumption) = [*N* (tdTomato^+^) / *N* (DAPI^+^)] × [*N* (c-Fos^+^) / *N* (DAPI^+^)]. To quantify activation and reactivation of rabies-defined input neurons in TRAP2::H2B-GFP mice (where H2B-GFP marks TRAP-captured activity and c-Fos marks reactivation), we computed: Activation of inputs = *N* (RV^+^H2B-GFP^+^) / *N* (RV^+^); Chance = *N* (H2B-GFP^+^) / *N* (DAPI^+^); Reactivation of inputs = *N* (RV^+^H2B-GFP^+^c-Fos^+^) / *N* (RV^+^); Chance = *N* (H2B-GFP^+^c-Fos^+^) / *N* (DAPI^+^). These definitions allowed direct comparisons between observed enrichment among rabies-labeled inputs and the corresponding chance levels expected from the regional cell population.

For synapse imaging, PSD-95-stained sections were imaged on an Olympus FV3000 confocal microscope using a 60× oil objective. Z-stacks were acquired at 1024 × 1024 pixels with 0.5 μm step size and 2-frame averaging, using identical acquisition settings across samples. For each slice, one dorsolateral striatal FOV at a matched anatomical location was selected; 3–4 FOVs per mouse were analyzed. In Imaris (v9.6.0), presynaptic boutons were segmented as synaptophysin-EGFP (M2) or synaptophysin-mTagBFP2 (PF) “spots”, TRAPed neurons were segmented as tdTomato^+^ surfaces (with soma/dendrite compartmentation when required), and PSD-95 puncta on tdTomato^+^ surfaces were segmented as postsynaptic “spots”. Putative synapses were defined by spatial co-localization of presynaptic and PSD-95 puncta on tdTomato^+^ surfaces using a centroid-to-centroid distance threshold tied to measured presynaptic spot size within each FOV. Synapse density was computed as synapse number normalized to tdTomato^+^ surface area, and coupling efficiency was computed as synapse number divided by total presynaptic boutons for each pathway within an FOV; FOV measures were averaged per mouse.

For nearest-neighbor analyses, synapse centroid coordinates were exported from Imaris and analyzed in MATLAB (R2022b). The nearest-neighbor distance (NND) was computed for each synapse as the Euclidean distance to its closest neighboring synapse within the FOV. Null distributions were generated by randomly reassigning synapse positions among available PSD-95(tdTomato^+^) puncta coordinates within each FOV (1,000 shuffles), preserving synapse count. Observed NND cumulative distributions were compared to shuffle-derived mean ± 95% confidence intervals; distributions were additionally expressed as Z-scored relative to shuffles for direct group comparison. The nearest-neighbor index (NNI) was calculated per FOV as mean observed NND divided by mean shuffled NND, and then averaged per mouse.

### Blinding, randomization, and exclusion criteria

Animals were randomly assigned to experimental groups when feasible, including levodopa versus withdrawal and virus or shRNA versus control comparisons, with cohort sizes balanced across experimental batches. Investigators were blinded to group identity during AIM scoring, offline behavioral quantification, and image quantification in ImageJ, Imaris, or MATLAB whenever blinding was compatible with the experimental design. Blinding was not feasible for some optogenetic experiments because implant location or stimulation configuration was apparent during testing.

Pre-established exclusion criteria were applied across experiments. Animals were excluded if histological verification showed off-target or insufficient viral expression, misplaced optical fibers, or inadequate dopamine depletion based on TH immunoreactivity. For ensemble-related behavioral experiments, animals were required to meet the AIM inclusion threshold described above after levodopa priming.^23^ For electrophysiology, cells were excluded if access resistance exceeded 30 MΩ or became unstable. For imaging-based contact analyses, FOVs were excluded if tissue damage, signal saturation, registration errors, or segmentation artifacts prevented reliable puncta detection.

### Statistics and reproducibility

Data are presented as mean ± SEM unless otherwise indicated. Sample sizes were not predetermined by formal power analysis but are comparable to prior studies using similar approaches. Statistical analyses were performed in GraphPad Prism (v8.0.1) and MATLAB (R2022b). Two-group comparisons used two-tailed unpaired or paired *t* tests when assumptions for parametric tests were met; otherwise, Mann–Whitney U tests or Wilcoxon matched-pairs signed rank tests were used. Multi-group comparisons used one-way or two-way ANOVA with appropriate post hoc tests as specified in figure legends. Categorical data were analyzed using Fisher’s exact test where appropriate. Significance thresholds were set at *p* < 0.05 (\**p* < 0.05, \*\**p* < 0.01, \*\*\**p* < 0.001, \*\*\*\**p* < 0.0001).

### Ethics statement

All animal experiments were reviewed and approved by the Institutional Animal Care and Use Committee of the Department of Laboratory Animal Science at Fudan University (Approval No. 2023-NZH-09JZS). All procedures were performed in accordance with institutional guidelines and ARRIVE guidelines.

## Results

### A reproducible striatal ensemble tracks and contributes to levodopa-induced dyskinesia

To identify circuit mechanisms that drive or restrain levodopa-induced dyskinesia (LID), we first established a unilateral 6-hydroxydopamine (6-OHDA) mouse model that reliably expressed dyskinesia after repeated levodopa exposure.^23,24^ Wild-type mice received 6-OHDA injections into the medial forebrain bundle (MFB), resulting in a profound loss of tyrosine hydroxylase (TH)-positive dopaminergic terminals in the ipsilateral striatum (Fig. S1A and B), together with the expected parkinsonian motor impairments (Fig. S1C–G). Repeated levodopa administration then elicited robust, time-locked dyskinesia that peaked ∼20–60 min after dosing and persisted for ∼2 h (Fig. S1H). LID was quantified using an established abnormal involuntary movement (AIM) framework^23,24^ that scores axial, limb, and orolingual dyskinesia on both temporal and amplitude scales (Fig. S1I).

The striatum integrates convergent long-range inputs that shape medium spiny neuron (MSN) activity and motor output.^27–31^ To gain access to the striatal neurons repeatedly engaged during LID, and thereby define a postsynaptic substrate for circuit dissection, we used FosTRAP^32,33^ in TRAP2::Ai14 mice to permanently label neurons engaged during a dyskinesia episode (Fig. S2). When dyskinesia was re-elicited in a subsequent levodopa session, a large fraction of TRAPed neurons was reactivated, yielding a reactivation rate well above chance (Fig. S2C–F), consistent with a stable striatal ensemble reproducibly recruited across dyskinesia bouts.^17^ TRAPed neurons broadly followed the spatial pattern of c-Fos expression, with enrichment in rostral and intermediate striatum (Fig. S2G) and in lateral sensorimotor-associated territories along the mediolateral axis (Fig. S2H and I).

We next asked whether this ensemble tracked dyskinesia expression in real time. Fiber photometry in TRAP2::Ai162 (TIT2L-GCaMP6s-ICL-tTA2) mice^34^ showed that TRAPed ensemble Ca^2+^ signals rose around dyskinesia onset and returned toward baseline as dyskinesia resolved, with elevated event frequency during the dyskinetic period relative to pre- and post-LID intervals (Fig. S2J–R). To test whether this population contributed to ongoing dyskinesia in our preparation, we optogenetically inhibited TRAPed striatal neurons. Inhibition acutely reduced LID, as reflected by a lower global AIM score relative to controls (Fig. S3). Together, these analyses establish FosTRAP-tagged striatal neurons as a reproducibly reactivated, LID-coupled, and behaviorally relevant ensemble suitable for upstream circuit analysis.

### M2 and PF are dominant afferents with opposite effects on dyskinesia

We then asked which long-range afferents are positioned to recruit the dyskinesia-linked striatal ensemble. We combined FosTRAP with Cre-dependent monosynaptic rabies tracing to label presynaptic neurons providing direct input to striatal TRAPed starter cells (Fig. 1A). Whole-brain quantification showed that ensemble inputs were dominated by cortex and thalamus, with additional contributions from basal ganglia nuclei including the globus pallidus (Fig. 1B and C). Within cortex, secondary motor cortex (M2) emerged as a prominent source of labeled afferents and contributed a larger input fraction than primary motor cortex (M1) in this tracing dataset (Fig. 1C). Within thalamus, intralaminar nuclei accounted for the majority of thalamic inputs,^35^ and the parafascicular nucleus (PF) comprised the dominant intralaminar contributor (Fig. 1D). These anatomical data identified M2 and PF as principal cortical and thalamic entry points to the dyskinesia-linked striatal ensemble.

**Fig. 1:**
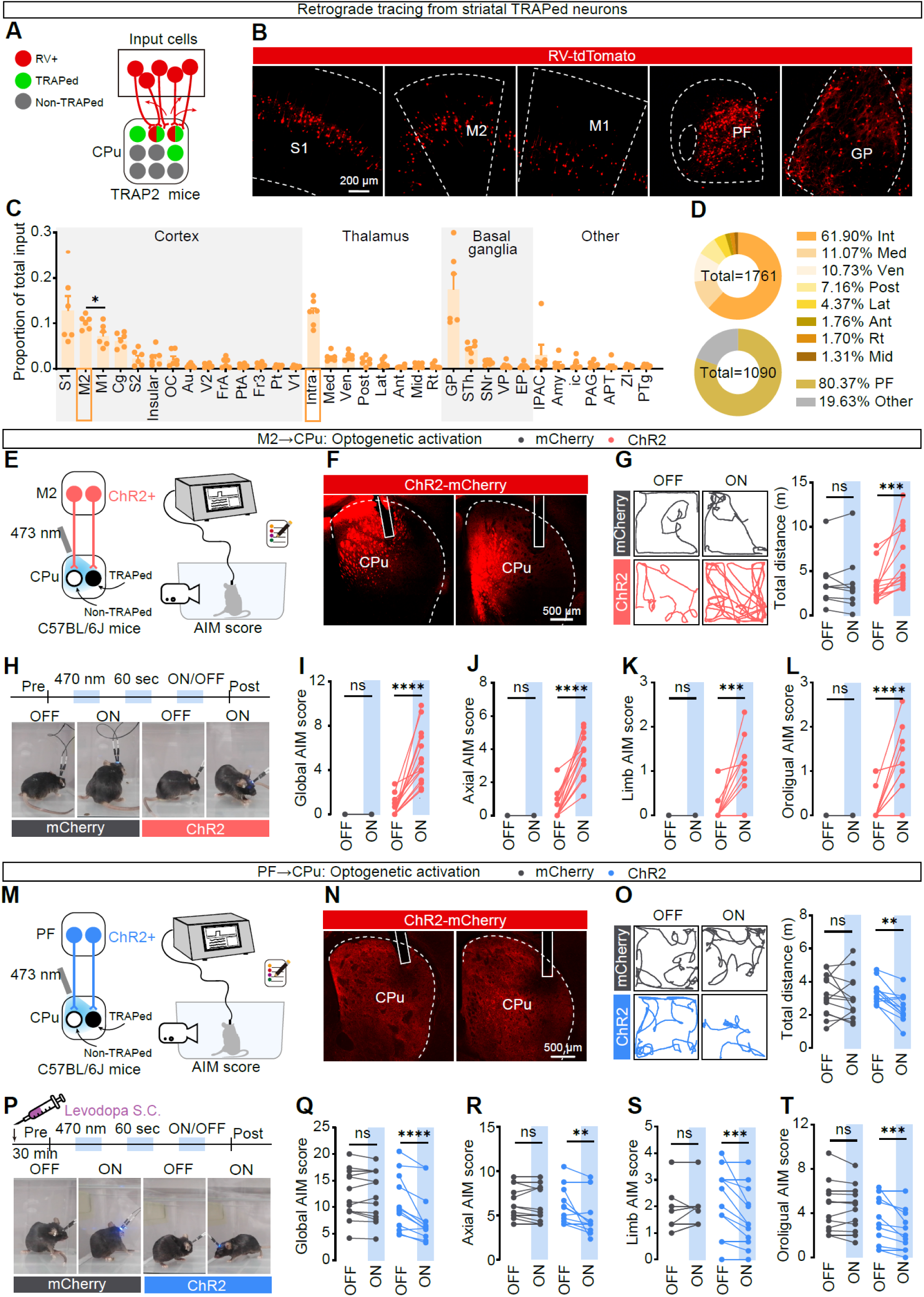
FosTRAP access to dyskinesia-linked striatal ensembles identifies dominant M2 and PF afferents with opponent behavioral effects. (A) Strategy for monosynaptic retrograde rabies tracing from FosTRAP-tagged striatal ensemble neurons. TRAPed starter cells in CPu were selectively complemented to permit infection by EnvA-pseudotyped ΔG rabies virus, thereby labeling presynaptic input neurons throughout the brain. (B) Representative RV-dsRed labeling in major upstream regions with prominent input to striatal TRAPed neurons, including primary somatosensory cortex (S1), secondary motor cortex (M2), primary motor cortex (M1), parafascicular thalamus (PF), and globus pallidus (GP). (C) Whole-brain quantification of extra-striatal input fractions to striatal TRAPed neurons. Brain regions are grouped by anatomical class and ordered by descending input strength within each class. M2 contributed a larger cortical input fraction than M1 (*N* = 6 mice; two-sided paired Student’s *t* test, \**p* = 0.0339). (D) Composition of thalamic inputs to striatal TRAPed neurons. Top, contribution of thalamic nuclear groups to total thalamic inputs (*n* = 1,761 traced thalamic neurons, pooled from *N* = 6 mice). Bottom, PF accounts for the majority of intralaminar inputs (*n* = 1,090 traced intralaminar neurons). Abbreviations: S1, primary somatosensory cortex; S2, secondary somatosensory cortex; M1, primary motor cortex; M2, secondary motor cortex; Cg, cingulate cortex; OC, orbital cortex; Au, auditory cortex; V1, primary visual cortex; V2, secondary visual cortex; FrA, frontal association cortex; PtA, parietal association cortex; Fr3, frontal cortex area 3; Pt, parietal cortex; PF, parafascicular thalamic nucleus; Int, intralaminar thalamic nuclei; Med, medial thalamic group; Ven, ventral thalamic group; Post, posterior thalamic group; Lat, lateral thalamic group; Ant, anterior thalamic group; Mid, midline thalamic group; Rt, reticular thalamic nucleus; GP, globus pallidus; STh, subthalamic nucleus; SNr, substantia nigra pars reticulata; VP, ventral pallidum; EP, entopeduncular nucleus; IPAC, interstitial nucleus of the posterior limb of the anterior commissure; Amy, amygdala; ic, internal capsule; PAG, periaqueductal gray; APT, anterior pretectal nucleus; ZI, zona incerta; PTg, pedunculotegmental nucleus. (E, M) Strategy for projection-wide optogenetic activation of M2 (E) or PF (M) terminals in dorsal striatum of 6-HDA–lesioned, levodopa-primed mice. ChR2-mCherry (or mCherry control) was expressed in ipsilateral M2 (E) or PF (M) and fibers were implanted over dorsolateral CPu for terminal stimulation during behavioral assays. (F, N) Representative ChR2-mCherry expression and optical fiber placements in CPu. (G, O) Open-field locomotion in the absence of levodopa during alternating light OFF/ON epochs. Left, representative 1-min center-point trajectories; right, total distance traveled. (G) M2, F_(1, 21)_ = 9.859, \*\**p* = 0.0049. mCherry, *N* = 9, ns, *p* = 0.9782; ChR2, *N* = 14, \*\*\**p* = 0.0002. (O) PF, F_(1, 23)_ = 4.268, ns, *p* = 0.0503; mCherry, *N* = 13, ns, *p* = 0.8089; ChR2, *N* = 12, \*\**p* = 0.0046. Two-way ANOVA followed by Sidak’s post hoc test. (H, P) Stimulation paradigm for AIM scoring (60-s OFF/ON epochs) and representative frames illustrating the emergence of dyskinesia-like abnormal movements during of M2→CPu (H) stimulation without levodopa and the alleviation of LID during PF→CPu (P) stimulation with levodopa. (I–L, Q–T) AIM score during OFF versus ON epochs for M2 and PF terminal stimulation (I and Q, global AIM score; J and R, axial; K and S, limb; L and T, orolingual). (I–L) M2, mCherry, *N* = 9; ChR2, *N* = 14. (I) F_(1, 21)_ = 27.79, \*\*\*\**p* < 0.0001; mCherry, ns, *p* > 0.9999; ChR2, \*\*\*\**p* < 0.0001. (J) F_(1, 21)_ = 61.50, \*\*\*\**p* < 0.0001; mCherry, ns, *p* > 0.9999; ChR2, \*\*\*\**p* < 0.0001. (K) F_(1, 21)_ = 6.790, \**p* = 0.0165; mCherry, ns, *p* > 0.9999; ChR2, \*\*\**p* = 0.0009. (L) M2, F_(1, 21)_ = 11.93, \*\**p* = 0.0024; mCherry, ns, *p* > 0.9999; ChR2, \*\*\*\**p* < 0.0001. (Q–T) PF, mCherry, *N* = 13; ChR2, *N* = 12. (Q) F_(1, 23)_ = 11.75, \*\**p* = 0.0023; mCherry, ns, *p* = 0.9177; ChR2, \*\*\*\**p* < 0.0001. (R) F_(1, 23)_ = 5.520, \**p* = 0.0278; mCherry, ns, *p* = 0.9031; ChR2, \*\**p* = 0.0027. (S) F_(1, 23)_ = 12.91, \*\**p* = 0.0015; mCherry, ns, *p* = 0.7419; ChR2, \*\*\**p* = 0.0005. (T) F_(1,23)_ = 5.692, \**p* = 0.0257; mCherry, ns, *p* = 0.8117; ChR2, \*\**p* = 0.0016. Two-way ANOVA followed by Sidak’s post hoc test. *n*, neurons; *N*, animals. Data are mean ± SEM.

Having identified M2 and PF as major monosynaptic afferents to the ensemble, we next tested whether these pathways exert distinct causal effects on dyskinesia expression *in vivo*. We expressed ChR2 in neurons in M2 or PF and implanted optical fibers over the dorsolateral striatum (caudate-putamen, CPu) of dyskinetic mice to stimulate M2→CPu or PF→CPu axon terminals (Fig. 1E, F, M, and N).

In dyskinetic mice tested during the inter-dose interval, when behavior resembled an OFF-like parkinsonian state, optogenetic stimulation of M2 terminals in CPu increased open-field locomotion (Figs. 1G, S4A and B), consistent with the pro-kinetic influence of corticostriatal drive.^36^ Notably, the same M2→CPu stimulation was sufficient to evoke AIMs in the absence of acute levodopa (Fig. 1H). AIM scoring revealed a robust increase in global dyskinesia score (Figs. 1I and S4C), with axial, limb, and orolingual components all contributing to the response (Figs. 1J–L and S4D–F). These stimulation-locked AIMs were reproducible across repeated OFF/ON epochs (Fig. S4D–F), indicating that transient enhancement of M2 corticostriatal drive can acutely trigger dyskinesia-like motor patterns.

PF projections exerted the opposite effect. When PF terminals in CPu were stimulated during the levodopa-free interval, mice showed reduced open-field locomotion (Figs. 1M–O and S5A–G), an effect that was also observed with chemogenetic activation of PF→CPu (Fig. S5H–L). We therefore asked whether PF engagement could suppress dyskinesia once it was already expressed. During ongoing levodopa-evoked dyskinesia, PF→CPu stimulation rapidly attenuated AIMs, often interrupting otherwise continuous abnormal movements (Fig. 1P). Across animals, PF→CPu activation reduced global AIM score as well as axial, limb, and orolingual subscores (Figs. 1Q–T and S5C–G). Repeated chemogenetic activation of PF→CPu also produced cumulative suppression of dyskinesia across days, with particularly prominent effects on axial AIMs (Fig. S5M–P). Thus, M2 and PF represent dominant afferent pathways with opposite behavioral effects: M2 acts as a pro-dyskinetic cortical drive, whereas PF acts as an anti-dyskinetic thalamostriatal brake.

### 3D behavioral mapping reveals opposite M2 and PF effects on dyskinetic state occupancy

AIM scoring provides a robust measure of dyskinesia severity, but it does not resolve how circuit perturbations reorganize the broader behavioral repertoire. We therefore asked whether M2 and PF manipulations simply changed movement amount or instead shifted animals toward or away from dyskinetic behavioral states. To address this, we applied a multi-view 3D pose-based behavioral mapping framework (BehaviorAtlas)^25^ to quantify optogenetically evoked motor patterns in an observer-independent manner (Fig. 2A). Four synchronized cameras captured open-field behavior, 16 body landmarks were tracked to reconstruct a 3D skeleton, and continuous trajectories were segmented into ∼1 s bouts. Unsupervised clustering yielded a library of recurring movement motifs, which we curated into 17 stereotyped behaviors grouped into dyskinesia, locomotion, exploration, and rest (Fig. 2B). Because fine orolingual movements are not reliably resolved by this pose representation, this analysis primarily captures axial and limb aspects of dyskinesia, whereas orolingual AIMs were quantified separately using standard scoring.

**Fig. 2:**
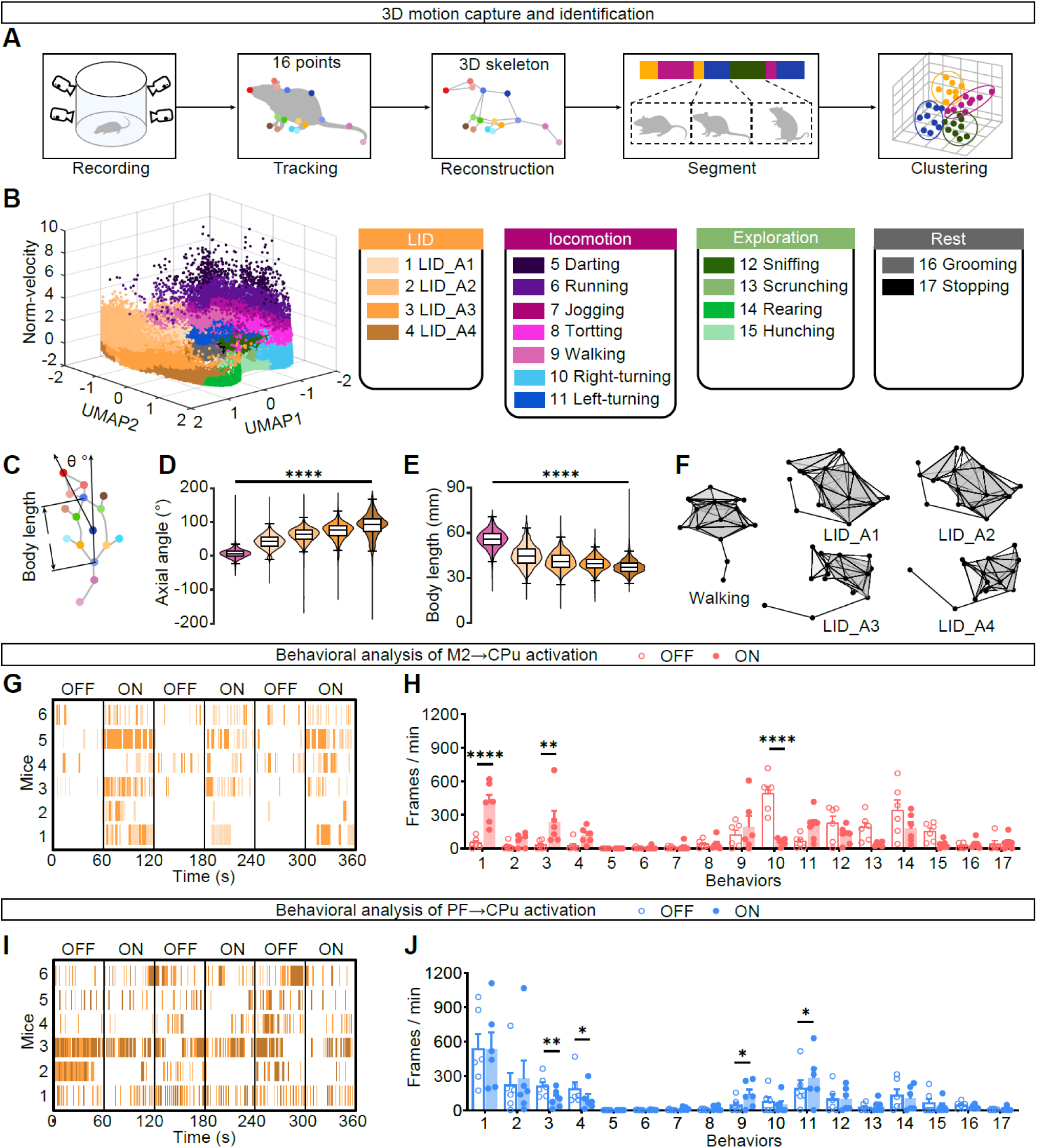
3D kinematic clustering shows that M2 and PF oppositely shift behavioral state occupancy between dyskinetic and non-dyskinetic regimes. (A) Schematic of the 3D multi-view motion-capture pipeline (BehaviorAtlas). Freely moving mice were recorded simultaneously form four calibrated viewpoints, 16 body landmarks were tracked, and a 3D skeleton was reconstructed. Continuous pose trajectories were segmented into short bouts and clustered in an unsupervised manner to define movement phenotypes. (B) Low-dimensional embedding of all segmented bouts pooled across recordings (axes: UMAP1, UMAP2, and normalized velocity). Each point denotes one bout, colored by the final curated set of 17 phenotypes. Phenotypes were grouped into four categories (LID, locomotion, exploration, and rest; key at right), including four dyskinesia-related classes (LID_A1–LID_A4). (C) Definition of kinematic features used to validate LID-related phenotypes. Body length was computed from the neck to tail base projection, and axial angle (θ) was defined from the body axis (tail base–back–nose). (D, E) Distributions of axial angle (D) and body length (E) across walking and LID_A1–LID_A4 bouts, showing progressively increased axial deviation and reduced body length with higher LID classes. (D) F_(4, 138194)_ = 40363.25, \*\*\*\**p* < 0.0001. (E) F_(4, 138194)_ = 25801.61, \*\*\*\**p* < 0.0001. Walking, *n* =16172 bouts; LID_A1, *n* = 52,225; LID_A2, *n* = 31,260; LID_A3, *n* = 19,576; LID_A4, *n* = 18,971; *N* = 24. One-way ANOVA with Tukey’s multiple comparisons. (F) Representative average skeleton templates for walking and LID_A1–LID_A4 generated by time-averaging landmark configurations within each phenotype. (G–J) Behavioral consequences of M2→CPu (G, H) or PF→CPu (I, J) activation quantified by 3D clustering. (G, I) Ethograms from individual ChR2 mice across alternating laser OFF/ON epochs (three cycles; 60 s per epoch) showing the temporal occurrence of LID phenotypes that changed significantly with stimulation. (H, J) Quantification of phenotype occupancy (frames/min) during OFF versus ON epochs across all 17 behaviors (key in B). (H) M2, F_(16, 85)_ = 11.02, \*\*\*\**p* < 0.0001; Behavior 1, \*\*\*\**p* < 0.0001; Behavior 3, \**p* = 0.0283; Behavior 10, \*\*\*\**p* < 0.0001. (J) PF, F_(16, 85)_ = 3.720, \*\*\*\**p* < 0.0001; Behavior 3, \*\**p* = 0.0016; Behavior 4, \**p* = 0.0283; Behavior 9, \**p* = 0.0153; Behavior 11, \**p* = 0.0208. OFF/ON measured within the same mice. *N* = 6. Two-way ANOVA with Sidak’s post hoc test. *n*, bouts; *N*, animals. All data presented as mean ± SEM or box-and-whisker plots.

Within the low-dimensional behavioral space, dyskinetic bouts formed a distinct region that could be stratified into four LID-related phenotypes (LID_A1–A4), aligned with increasing axial and limb abnormality (Fig. 2B). Quantitative kinematic features validated this ordering: higher LID classes showed progressively greater axial torsion, elevated trunk and forepaw positions, and reduced body length, consistent with increasingly abnormal dyskinetic postures (Figs. 2C–F and S6A–D). These analyses established that the 3D pipeline distinguishes graded dyskinesia-like behaviors from normal locomotor and exploratory actions.

We then generated ethograms for each animal and compared behavioral motif occupancy during laser OFF versus ON epochs. In levodopa-free sessions, M2→CPu stimulation produced a rapid redistribution of behavior into dyskinesia-related motifs (Fig. 2G and H). LID_A1 and LID_A3 bouts emerged prominently during stimulation and were rarely observed during OFF epochs, consistent with the AIM scoring results. At the same time, parkinsonian ipsilateral turning and exploratory behaviors were reduced (Fig. 2H), indicating that M2 drive does not merely increase movement amount but shifts the repertoire toward a dyskinesia-dominated state. Control mice expressing mCherry showed no comparable state shift (Fig. S6E and F).

In contrast, during peak LID, PF→CPu activation shifted the behavioral repertoire away from the most severe dyskinetic motifs (Fig. 2I and J). High-amplitude LID behaviors (LID_A3 and LID_A4) were selectively reduced, whereas milder dyskinesia motifs were comparatively spared. Importantly, this reduction was accompanied by increased expression of non-dyskinetic behaviors, including locomotor modules such as walking and turning, rather than uniform movement suppression (Fig. 2J). Thus, PF activation shifted ongoing motor output away from severe dyskinesia and toward a more physiological behavioral repertoire. Together, 3D behavioral mapping shows that M2 and PF act as opponent controllers of behavioral state occupancy: M2 pushes animals toward a dyskinetic state, whereas PF reduces occupancy of dyskinetic states and favors return toward non-dyskinetic behavior.

### Chronic levodopa preserves pathway-to-ensemble coupling despite terminal loss

Given the opposite behavioral effects of M2 and PF activation (Fig. 1) and the state shifts revealed by 3D behavior mapping (Fig. 2), we next asked whether chronic levodopa remodels the anatomical interface between these afferents and the dyskinesia-linked striatal ensemble. To quantify pathway-specific structural coupling, we labeled M2 and PF presynaptic boutons with synaptophysin-tagged EGFP or BFP, respectively, in TRAP2::Ai14 mice, and immunostained PSD95 to mark excitatory postsynaptic sites (Fig. 3A and B). Putative M2→ensemble and PF→ensemble contacts were operationally defined as presynaptic synaptophysin puncta apposed to PSD95 puncta on tdTomato-labeled ensemble neurites (Figs. 3A and S7A). Levodopa continuation and withdrawal cohorts were matched for dyskinesia severity at the time of labeling and showed comparable local ensemble neuron density within imaged fields of view (Figs. 3C, H and S7B–E).

**Fig. 3:**
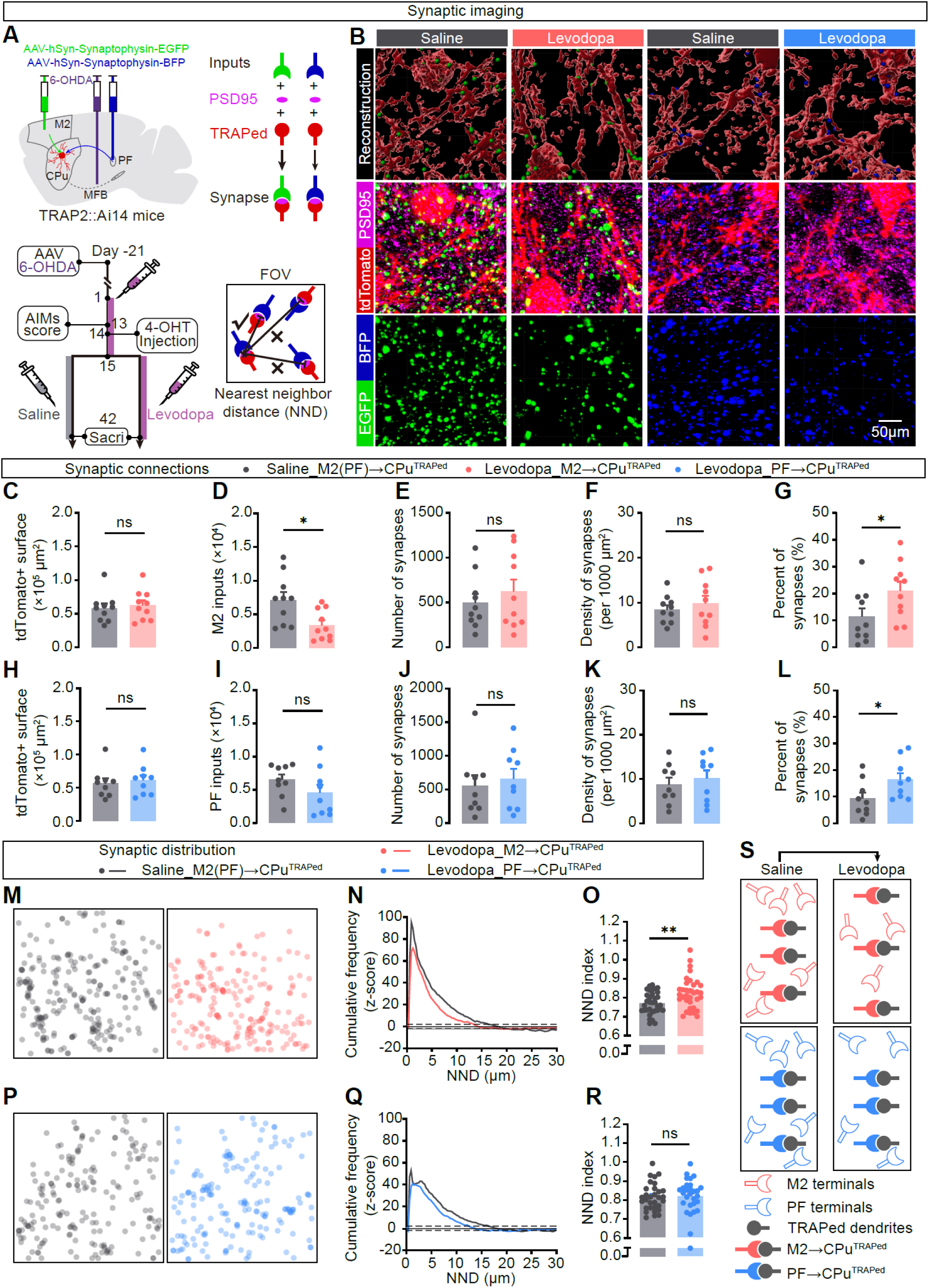
Chronic levodopa preserves pathway-to-ensemble contacts despite loss of presynaptic terminals and selectively disperses M2 contact organization. (A) Experimental design and analysis pipeline for dual-color synapse mapping onto FosTRAP-tagged ensemble neurons. In TRAP2::Ai14 mice, synaptophysin–EGFP and synaptophysin–BFP were expressed in M2 and PF axon terminals, respectively, and dyskinesia-activated striatal ensemble neurons were labeled by tdTomato after TRAP induction. Excitatory postsynaptic densities were visualized by PSD95 immunostaining. Putative pathway-specific synaptic contacts onto TRAPed neurons were defined by the apposition of a presynaptic synaptophysin punctum (EGFP or BFP) with a PSD95 punctum within the tdTomato^+^ neuronal surface. For spatial organization analysis, the nearest-neighbor distance (NND) between synapse centroids was computed within each field of view (FOV) and compared to shuffled synapse maps. (B) Representative confocal images and corresponding 3D reconstructions from dorsolateral striatum in saline- and chronic levodopa-treated mice. TRAPed neurons (tdTomato, red), PSD95 (magenta), and presynaptic terminals from M2 (synaptophysin–EGFP, green) or PF (synaptophysin– BFP, blue) are shown. (C–L) Quantification of M2→CPu^TRAPed^ or PF→CPu^TRAPed^ structural coupling. tdTomato^+^ surface area was unchanged by chronic levodopa (C, *p* = 0.6599; H, *p* = 0.6997; two-tailed unpaired Student’s *t* test). The total number of M2 terminals (synaptophysin–EGFP puncta) in the sampled volume was reduced (D, \**p* = 0.0133; two-tailed unpaired Student’s *t* test), whereas PF terminal counts (synaptophysin–BFP puncta) did not differ significantly between groups (I, *p* = 0.1942; two-tailed unpaired Student’s *t* test). The number (E, *p* = 0.4682; two-tailed unpaired Student’s *t* test. J, *p* = 0.7304; Mann-Whitney U test) and density (normalized to tdTomato^+^ surface) (F, *p* = 0.4792; K, *p* = 0.5556; two-tailed unpaired Student’s *t* test) of M2 (E, F) or PF (J, K) contacts onto TRAPed neurons were maintained, resulting in an increased fraction of M2 (G, \**p* = 0.0473; two-tailed unpaired Student’s *t* test) or PF (L, \**p* = 0.0482; two-tailed unpaired Student’s *t* test) terminals contacting TRAPed neurons. (C–G) Saline, *N* = 10; Levodopa, *N* = 10. (H–L) Saline, *N* = 9; Levodopa, *N* = 9. (M–R) Spatial distribution of M2→CPu^TRAPed^ (M–O) of PF→CPu^TRAPed^ (P–R) contacts. (M, P) Representative synapse location maps from single FOVs in saline (left) and levodopa (right) groups. Synapses are represented by colored circles with set transparency so that the depth of the color represents the density of the distribution. (N, Q) Cumulative NND distributions plotted as Z scores relative to shuffled synapse maps; dashed line indicates the 95% confidence interval of the shuffled distribution. (N) M2→CPu^TRAPed^ synapses. Saline, *n* = 17,356 synapses, *N* = 10; Levodopa, *n* = 19,005 synapses, *N* = 10. \*\*\**p* = 0.003; Kolmogorov-Smirnov test. (Q) PF→CPu^TRAPed^ synapses. Saline, *n* = 16,453 synapses, *N* = 9; Levodopa, *n* = 18,153 synapses, *N* = 9. ns, *p* = 0.1399; Kolmogorov-Smirnov test. (O) Nearest-neighbor index (NNI; mean observed NND divided by mean shuffled NND) was increased after chronic levodopa in M2→CPu^TRAPed^ synapses, consistent with reduced clustering (greater dispersion) of M2 contacts. Saline, *n* = 34, *N* = 10; Levodopa, *n* = 34, *N* = 10. \*\**p* = 0.0027. (R) Meanwhile, NNI was unchanged by chronic levodopa in PF→CPu^TRAPed^ synapses, indicating preserved PF contact organization. Saline, *n* = 31, *N* = 9; Levodopa, *n* = 29, *N* = 9. ns, *p* = 0.7809. Two-tailed unpaired Student’s *t* test. (S) Schematic summary: chronic levodopa decreases overall presynaptic terminal availability but preserves pathway-specific contacts onto LID-linked ensemble neurons, increasing the fraction of surviving M2 and PF terminals that couple to TRAPed targets; spatial dispersion is selectively observed for M2 contacts. *n*, slices; *N*, animals. Data are mean ± SEM.

Chronic levodopa exposure produced a reduction in overall corticostriatal presynaptic bouton abundance, with significantly fewer M2 synaptophysin puncta in the striatum (Fig. 3D) and a trend toward reduced PF puncta (Fig. 3I). However, the absolute number and density of putative contacts onto ensemble neurons were preserved for both pathways: neither M2→ensemble nor PF→ensemble contact counts differed between chronic levodopa and withdrawal conditions (Fig. 3E, F, J, and K). Consequently, the fraction of remaining presynaptic boutons contacting ensemble neurites increased for both inputs (Fig. 3G and L), indicating enhanced pathway-to-ensemble coupling despite contraction of the broader terminal fields. These data suggest that chronic levodopa does not globally increase corticostriatal and thalamostriatal innervation, but instead preserves effective synaptic access to dyskinesia-linked ensemble neurons.

Because the spatial arrangement of excitatory inputs can shape dendritic integration, we next examined whether chronic levodopa also remodels the spatial organization of M2 and PF contacts onto ensemble neurons. Using the same synapse-imaging dataset, we quantified nearest-neighbor distances (NNDs) among input-defined contacts on TRAPed neurons and compared the observed distributions with within-field shuffled controls (Fig. S7F–I).

Across pathways and conditions, NND distributions were consistently left-shifted relative to shuffled controls, indicating non-random spatial organization of afferent contacts on ensemble neurons (Fig. S7F–I). To compare conditions directly, cumulative NND distributions were Z-scored relative to their shuffled counterparts.^37^ Chronic levodopa selectively weakened local clustering of M2 contacts: M2→ensemble contacts shifted toward larger NNDs, showed reduced deviation from shuffled distributions, and exhibited an increased nearest-neighbor index (NNI), consistent with a more spatially dispersed arrangement (Fig. 3M–O). In contrast, PF→ensemble contacts maintained stable NND distributions and NNI across conditions (Fig. 3P–R), indicating preserved spatial organization despite increased coupling efficiency. Thus, chronic levodopa preserves effective pathway-to-ensemble coupling for both inputs, but selectively disperses M2 contacts while leaving PF contact organization largely intact (Fig. 3S).

### M2 and PF impose distinct synaptic mechanisms on dyskinesia-linked ensembles

We next asked whether the structurally remodeled M2 and PF inputs are also functionally remodeled in dyskinetic animals. We performed whole-cell electrophysiological recordings in dorsolateral striatal slices from TRAP2::Ai14 mice, targeting tdTomato-labeled TRAPed ensemble neurons and neighboring non-TRAPed neurons for within-slice comparison.

At baseline, spontaneous excitatory drive was comparable between TRAPed and non-TRAPed neurons, as reflected by unchanged sEPSC frequency and amplitude (Fig. S8A–C). In contrast, TRAPed neurons showed a selective increase in sIPSC frequency without a change in amplitude (Fig. S8D–F), indicating stronger spontaneous inhibitory input onto ensemble neurons in the dyskinetic state, although the source of this inhibition was not resolved by spontaneous recordings alone.

To isolate pathway-specific responses, we expressed an excitatory opsin in M2 or PF and photostimulated their striatal terminals while recording optogenetically evoked synaptic currents (Fig. 4A and K). Both pathways displayed altered paired-pulse dynamics in TRAPed relative to non-TRAPed neurons, reflected by reduced paired-pulse ratios during brief stimulus trains (Fig. 4B, C, L, and M). We interpret these changes conservatively as altered short-term dynamics consistent with enhanced effective transmission during repeated activation, rather than as a definitive measure of presynaptic release probability. Despite this shared functional signature, the two pathways diverged in how ensemble neurons integrated their signals.

**Fig. 4:**
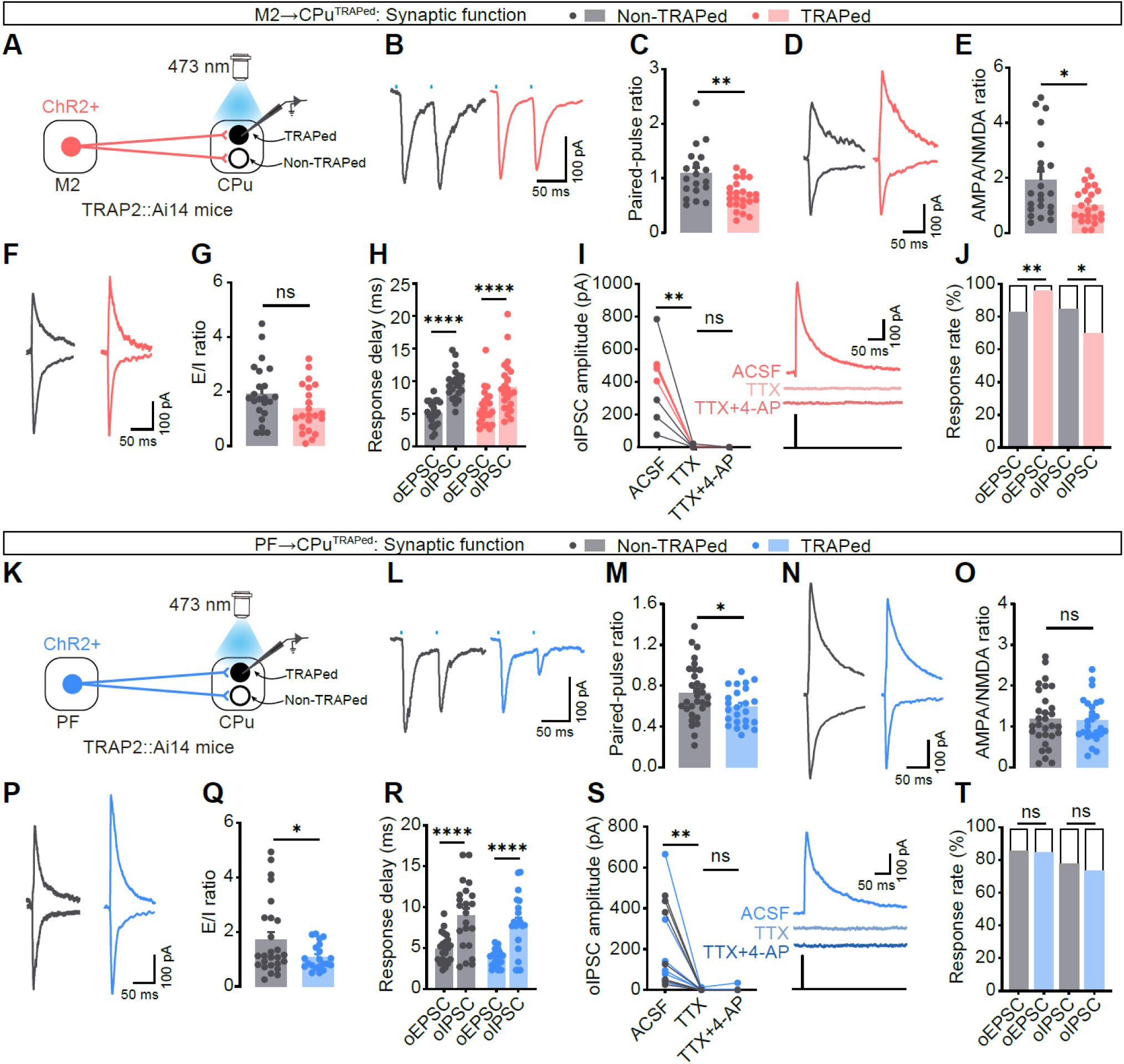
M2 and PF impose distinct synaptic integration rules on dyskinesia-linked ensemble neurons. (A, K) Recording configurations for pathway-isolated synaptic measurements in the dorsolateral striatum of TRAP2::Ai14 mice. Opsin-expressing axon terminals from M2 (A) or PF (K) were activated by blue light (473 nm) while whole-cell voltage-clamp recordings were obtained from neighboring tdTomato^+^ TRAPed and tdTomato^−^ non-TRAPed neurons. (B, C, L, M) M2-(B) or PF-evoked (L) paired-pulse oEPSCs and summary of paired-pulse ratio (PPR; 50 ms inter-stimulus interval; second/first peak amplitude) (C, M). (C) M2, nonTRAPed, *n* = 19, *N* = 8; TRAPed, *n* = 24, *N* = 8. \*\**p* = 0.0011. (M) PF, nonTRAPed, *n* = 31, *N* = 8 mice; TRAPed, *n* = 24, *N* = 8. \**p* = 0.0373. Two-tailed unpaired Student’s *t* test. (D, E, N, O) Representative M2-(D) or PF-evoked (E) AMPAR- and NMDAR-mediated currents and summary of AMPA/NMDA ratio (E, O). AMPAR component was measured as the oEPSC peak at −70 mV; NMDAR component was measured at +40 mV at 50 ms after stimulation. (E) M2, nonTRAPed, *n* = 22, *N* = 9; TRAPed, *n* = 24, *N* = 9. \**p* = 0.0329; Mann-Whitney U test. (O) PF, nonTRAPed, *n* = 30, *N* = 10; TRAPed, *n* = 26, *N* = 10. ns, *p* = 0.7772; two-tailed unpaired Student’s *t* test. (F, G, P, Q) Representative M2-(F) or PF-evoked (P) oEPSCs (−70 mV) and oIPSCs (0 mV) and evoked excitation/inhibition ratio (E/I; oEPSC peak/oIPSC peak) (Q, Q). (G) M2, nonTRAPed, *n* = 23, *N* = 7; TRAPed, *n* = 22, *N* = 7. ns, *p* = 0.066; two-tailed unpaired Student’s *t* test. (Q) PF, nonTRAPed, *n* = 23, *N* = 8; TRAPed, *n* = 21, *N* = 8. \**p* = 0.0216; Mann-Whitney U test. (H, R) Response delay (onset latency from stimulation) for M2-(H) or PF-evoked (R) oEPSCs and oIPSCs recorded in the same cells. (H) M2, nonTRAPed, *n* = 22, *N* = 7. \*\*\*\**p* < 0.0001; two-tailed paired Student’s *t* test. TRAPed, *n* = 22, *N* = 7. \*\*\*\**p* < 0.0001; Wilcoxon matched-pairs signed rank test. (R) PF, nonTRAPed, *n* = 22, *N* = 8, \*\*\*\**p* < 0.0001; TRAPed, *n* = 20 cells, *N* = 8 mice. Two-tailed paired Student’s *t* test. (I, S) Pharmacological test of the inhibitory component evoked by M2 (I) or PF (S) stimulation. oIPSCs were measured in ACSF, after TTX (1 μM), and after TTX + 4-AP (200 μM). Left, group data; right, example traces. (I) M2, *n* = 8 (non-TRAPed = 4, TRAPed = 4), *N* = 2. ASCF vs. TTX, \*\**p* = 0.0029; TTX vs. TTX + 4-AP, ns, *p* = 0.4838. One-way RM ANOVA followed by Tukey’s post hoc test. (S) PF, *n* = 13 (non-TRAPed = 7, TRAPed = 6), *N* = 3. ASCF vs. TTX, \*\**p* = 0.0064; TTX vs. TTX + 4-AP, ns, *p* = 0.7433. One-way RM ANOVA followed by Tukey’s post hoc test. (J, T) Response probability (percentage of recorded cells exhibiting detectable oEPSCs or oIPSCs) during M2 (J) or PF (T) terminal stimulation. (J) M2, oEPSC, *N* = 9, \*\**p* = 0.0046; oIPSC, *N* = 9; \**p* = 0.017. Fisher’s exact test. (T) PF, oEPSC, *N* = 10, ns, *p* > 0.9999; oIPSC, *N* = 10; ns, *p* = 0.6191. Fisher’s exact test. *n*, cells; *N*, animals. Data are as mean ± SEM.

At M2→ensemble synapses, optogenetically evoked EPSCs were relatively enriched for NMDAR-mediated currents, resulting in a lower AMPA/NMDA ratio in TRAPed neurons than in neighboring non-TRAPed neurons (Fig. 4D and E). By contrast, the overall evoked excitation-to-inhibition ratio was not detectably altered (Fig. 4F and G). Inhibitory currents recruited by M2 stimulation followed oEPSCs with longer onset latencies (Fig. 4H) and were abolished by TTX without recovery by 4-AP (Fig. 4I), consistent with polysynaptic feedforward inhibition rather than direct inhibitory input.^26^ Notably, response success shifted toward excitation, with increased oEPSC response probability and decreased oIPSC response probability in TRAPed neurons (Fig. 4J). Thus, chronic levodopa remodels M2 influence on ensemble neurons primarily by biasing glutamatergic signaling toward NMDAR-dependent excitation, rather than by producing a large shift in evoked excitation-to-inhibition balance.

PF→ensemble signaling followed a different rule. PF inputs did not exhibit NMDAR enrichment: the AMPA/NMDA ratio at PF→ensemble synapses was unchanged (Fig. 4N and O). Instead, PF stimulation recruited a disproportionately strong inhibitory component, reflected by a reduced oEPSC/oIPSC ratio in TRAPed neurons (Fig. 4P and Q), consistent with the elevated spontaneous inhibitory drive observed in ensemble neurons (Fig. S8F). Because PF projection neurons are predominantly glutamatergic,^38^ we tested whether PF-evoked oIPSCs arose through recruitment of local striatal inhibitory microcircuits. PF-evoked oIPSCs were delayed relative to PF-evoked oEPSCs (Fig. 4R) and were eliminated by TTX without rescue by 4-AP (Fig. 4S), indicating an action potential-dependent polysynaptic origin. Together, these recordings show that M2 and PF engage dyskinesia-linked ensemble neurons through distinct synaptic mechanisms: M2 inputs are biased toward NMDAR-dependent excitation, whereas PF inputs are routed through stronger polysynaptic inhibition that shifts net synaptic drive toward suppression.

### Circuit-restricted stimulation reveals an ensemble-external PF brake

Given the opposite behavioral effects of M2 and PF and their distinct synaptic mechanisms, we next asked whether the upstream neurons innervating the dyskinesia-linked ensemble are themselves preferentially engaged during dyskinesia. We combined activity tagging (TRAP2::H2B-GFP), monosynaptic rabies tracing from striatal ensemble neurons, and post hoc c-Fos mapping (Fig. 5A–C). H2B-GFP labeled neurons engaged during the TRAP window, rabies virus (RV-dsRed) labeled monosynaptic inputs to the ensemble, and c-Fos identified neurons reactivated during a subsequent dyskinesia episode. This strategy allowed us to quantify, within each upstream region, the fraction of ensemble-projecting neurons engaged during dyskinesia induction (RV^+^H2B-GFP^+^) or re-engaged during dyskinesia expression (RV^+^H2B-GFP^+^c-Fos^+^), each benchmarked against regional chance levels (Fig. 5D–I).

**Fig. 5:**
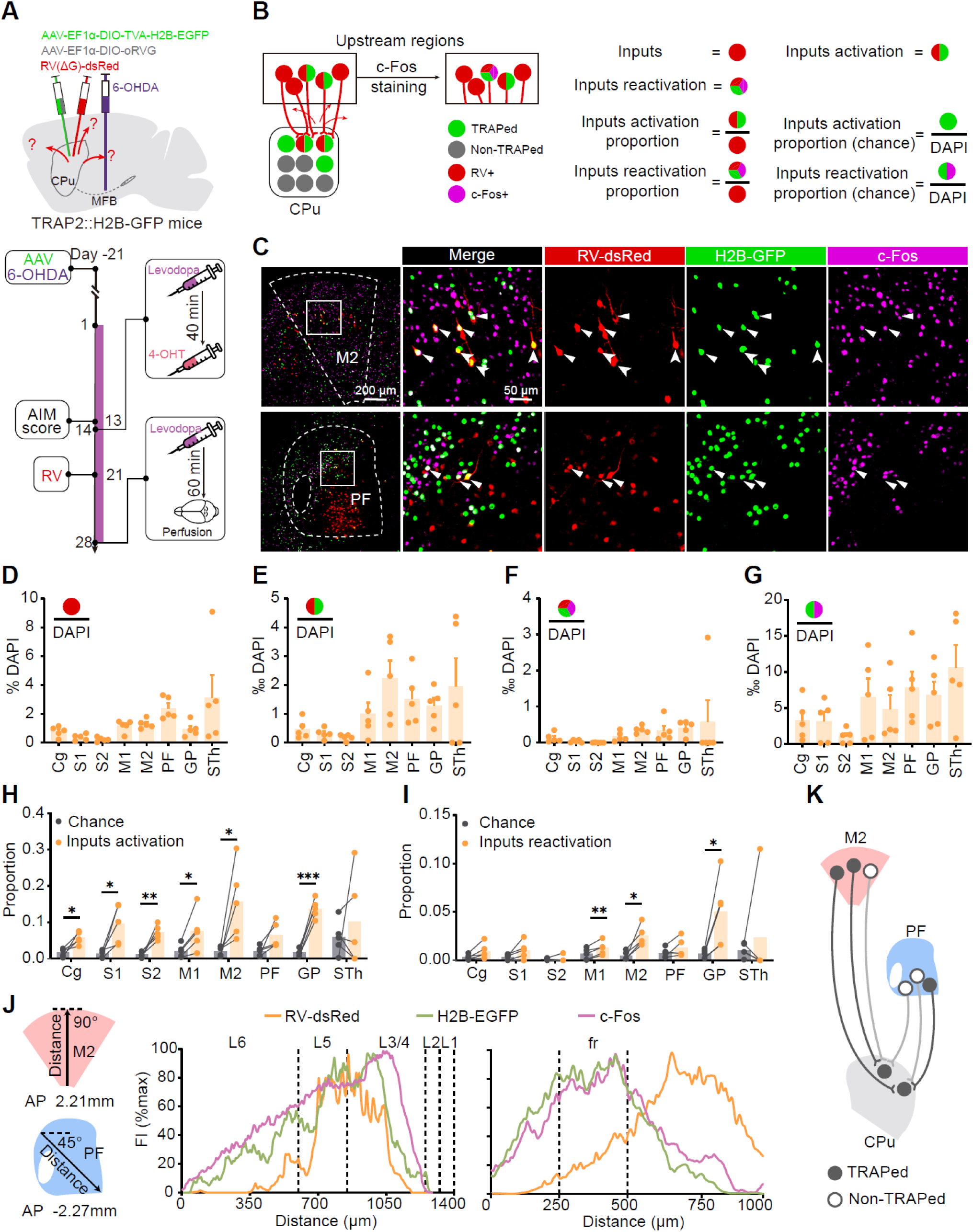
Dyskinesia-linked activity aligns with M2 ensemble connectivity but is dissociated from PF ensemble-projecting topology. (A) Experimental design to simultaneously label monosynaptic inputs to striatal TRAPed ensemble neurons and to read out their activation across LID episodes. In TRAP2::H2B-GFP mice, rabies helper AAVs were delivered to dorsal striatum, followed by EnvA pseudotyped ΔG rabies virus (RV-dsRed) to label upstream inputs. During a LID episode, 4-OHT was administered to TRAP (H2B-GFP) neurons activated in that episode; mice were challenged again with levodopa and processed for c-Fos immunostaining to mark reactivated neurons. (B) Operational definitions used for quantification. RV^+^ (dsRed^+^) neurons in upstream regions were classified as input neurons; “activated inputs” were RV^+^H2B-GFP^+^ neurons (inputs active during the TRAP episode); “reactivated inputs” were RV^+^H2B-GFP^+^c-Fos^+^ neurons (inputs active during both episodes). Input activation/reactivation fractions were compared with region matched chance levels estimated from population labeling rates (see Methods). (C) Representative images from M2 (top) and PF (bottom) showing rabies-labeled inputs (RV-dsRed, red), TRAP-labeled neurons (H2B-GFP, green), and reactivated neurons (c-Fos, magenta). Arrowheads indicate activated input neurons (RV^+^H2B-EGFP^+^); triangles indicate reactivated input neurons (RV^+^H2B-EGFP^+^c-Fos^+^). (D–G) Regional distribution of labeled populations in candidate input structures, quantified as percent of DAPI^+^ cells: (D) total rabies-labeled inputs (RV^+^), (E) activated inputs (RV^+^H2B-GFP^+^), (F) reactivated inputs (RV^+^H2B-GFP^+^C-Fos^+^), and (G) reactivated neurons in the regional population (H2B-GFP^+^c-Fos^+^). *N* = 5. (H, I) Enrichment of rabies-defined inputs above chance. For each region, the observed fraction of (H) activated inputs and (I) reactivated inputs was compared with the corresponding chance estimate within the same mouse. Ensemble projecting inputs in M2 were significantly enriched for both activation and reactivation, whereas PF inputs were not. *N* = 5. (H) Cg, \**p* = 0.0191; S1, \**p* = 0.0286; S2, \*\**p* = 0.0065; M1, \**p* = 0.0327; M2, \**p* = 0.0398; PF, *p* = 0.595; GP, \*\*\**p* = 0.0003; STh, ns, *p* = 0.5398. Two-tailed paired Student’s *t* test. (I) Cg, *p* = 0.068; S1, *p* = 0.0857; S2, *p* = 0.625; M1, \*\**p* = 0.0043; M2, \**p* = 0.0161; PF, *p* = 0.1862; GP, \**p* = 0.0376; STh, *p* = 0.625. S2 and STh, Wilcoxon matched-pairs signed rank test; others, two-tailed paired Student’s *t* test. (J) Spatial relationship between ensemble connectivity and LID-linked activity. Fluorescence intensity profiles for RV-dsRed (inputs), H2B-GFP (TRAP), and c-Fos across M2 (radial axis; AP +2.21 mm) and across PF (45° axis; AP −2.27 mm). Dashed lines mark cortical layer boundaries and the fasciculus retroflexus (fr). Intensities were normalized to the regional maximum within each section. *N* = 5. (K) Summary schematic illustrating preferential reactivation of ensemble projecting M2 neurons and the PF dissociation between the LID-reactivated population and the ensemble projecting population. *N*, animals. Data are mean ± SEM. Abbreviations: Cg, cingulate cortex; S1/S2, primary/secondary somatosensory cortex; M1/M2, primary/secondary motor cortex; PF, parafascicular thalamic nucleus; GP, globus pallidus; STh, subthalamic nucleus; CPu, caudate–putamen.

Across the input-rich network identified by rabies tracing—including motor cortices, PF, pallidum, and subthalamic nucleus (Fig. 5D–G and S9A)—ensemble-projecting neurons in M2 showed a selective bias toward activation and reactivation. The fraction of M2 inputs to the ensemble that were activity-tagged during dyskinesia induction exceeded chance, and this enrichment persisted among reactivated inputs during dyskinesia expression (Fig. 5H and I). By contrast, despite robust overall activity in PF during LID, as reflected by H2B-GFP and c-Fos labeling (Fig. 5E–G), PF neurons projecting monosynaptically to the striatal ensemble did not show enrichment above chance for activation or reactivation (Fig. 5H and I). Representative sections showed sparse overlap among RV-labeled PF inputs, H2B-GFP, and c-Fos (Fig. 5C). Thus, dyskinesia preferentially engages an ensemble-projecting cortical population in M2, whereas PF activity during dyskinesia does not preferentially map onto PF neurons that directly innervate the ensemble.

We then examined the spatial relationship between connectivity and activity within each upstream structure. In M2, RV-labeled ensemble-projecting neurons were concentrated in layers that overlapped extensively with both H2B-GFP-tagged and c-Fos^+^ populations (Fig. 5J), indicating that dyskinesia-linked cortical activity substantially coincides with M2 neurons projecting to the ensemble. In PF, however, the distributions were topographically segregated: RV-traced ensemble-projecting neurons were biased to lateral PF—the sector known to preferentially target lateral striatum—whereas H2B-GFP- and c-Fos-labeled neurons clustered more medially and centrally (Fig. 5J), consistent with the known organization of PF→striatum projections along this axis.^22,39^ These data reveal a topographic dissociation in PF whereby dyskinesia-linked activity is spatially segregated from the PF subpopulation that directly innervates the dyskinesia-linked striatal ensemble (Fig. 5K).

This dissociation suggested different circuit logic for M2 and PF. M2 appeared to engage a dyskinesia-linked corticostriatal subnetwork that converges onto ensemble neurons, whereas PF influence during dyskinesia might arise from a thalamic subpopulation that does not directly target the ensemble. To test this, we used an ensemble-restricted monosynaptic strategy to express ChR2 selectively in neurons that directly innervate striatal TRAPed neurons. In TRAP2 mice, Cre-dependent helper viruses (TVA and rabies glycoprotein) were first introduced into striatal starter cells, followed by EnvA-pseudotyped ΔG rabies encoding ChR2 to label monosynaptic inputs to the ensemble (Fig. S10A and B). Optical fibers were then implanted above the presynaptic cell-body regions (M2 or PF) to stimulate the ensemble-projecting subcircuits (Fig. 6).

**Fig. 6:**
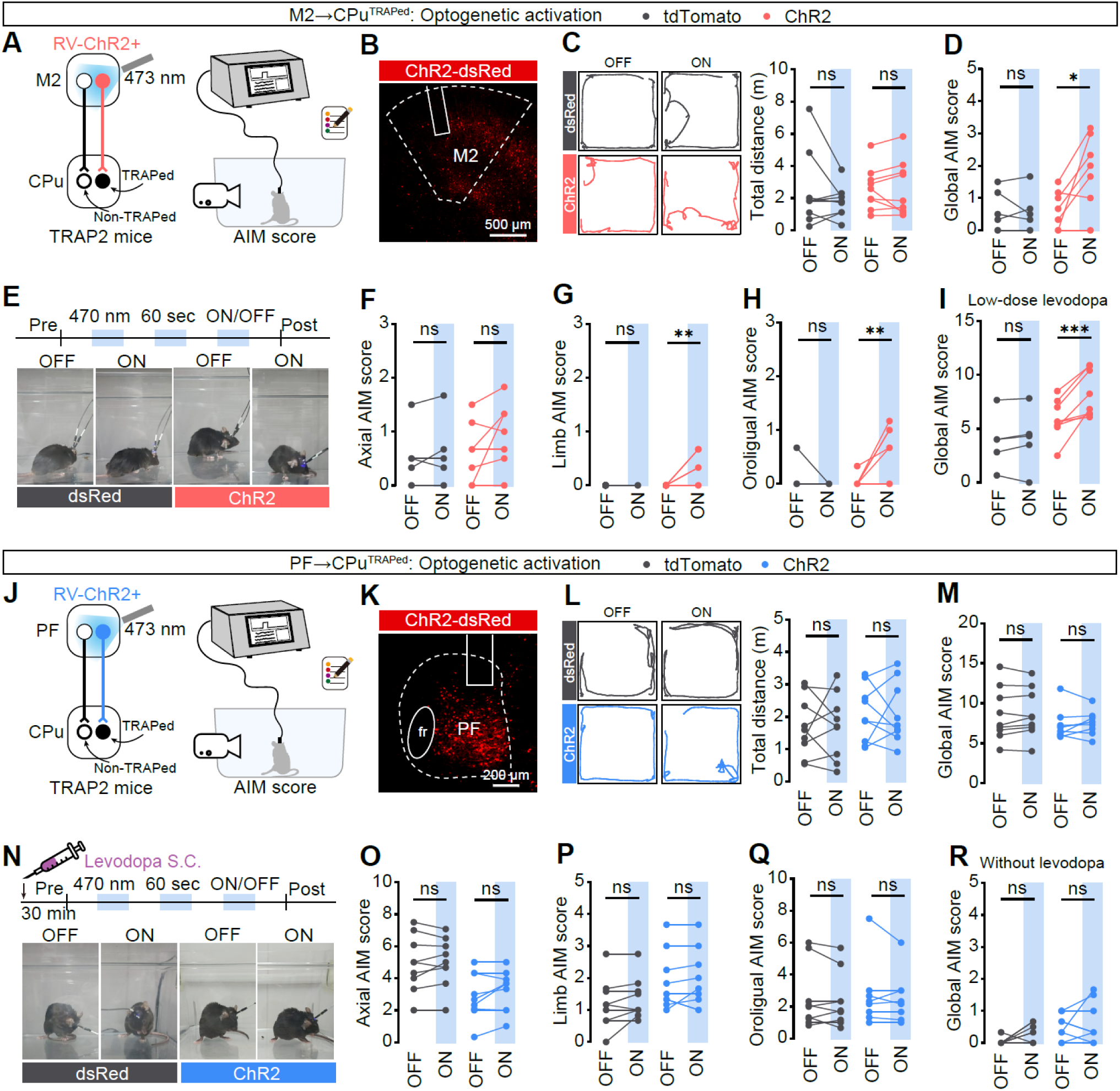
Circuit-restricted stimulation isolates an M2-to-ensemble driver and an ensemble-external PF brake. (A, J) Schematic of circuit-restricted optogenetic activation of the M2→CPu^TRAPed^ (A) or PF→CPu^TRAPed^ (J) pathway. Rabies viruses encoding ChR2–dsRed were injected into CPu of TRAP2 mice to retrograde infect presynaptic inputs and selectively express opsin in ensemble-projecting presynaptic neurons. Optic fibers were implanted over M2 or PF to photostimulate the rabies-labeled M2 or PF input population and quantify locomotion and AIMs. (B, K) Representative coronal section showing rabies-mediated ChR2-dsRed expression in M2 (b) or PF (k) and optical fiber placement. (C, L) Left, representative 1-min center-point trajectories during light OFF and ON epochs; right, total distance traveled in the open field during OFF versus ON epochs during M2→CPu^TRAPed^ or PF→CPu^TRAPed^ stimulation. (C) M2→CPu^TRAPed^, F_(1, 16)_ = 1.176, *p* = 0.2943; tdTomato, *p* = 0.2725; ChR2, *p* > 0.9999. (L) PF→CPu^TRAPed^, F_(1, 16)_ = 0.001421, *p* = 0.9704; tdTomato, *p* = 0.9995; ChR2, *p* = 0.9958. tdTomato, *N* = 9; ChR2, *N* = 9. Two-way ANOVA followed by Sidak’s post hoc test. (D, M) Global AIM score during circuit-restricted M2→CPu^TRAPed^ (D) or PF→CPu^TRAPed^ (M) stimulation. (D) M2→CPu^TRAPed^, F_(1, 16)_ = 7.120, \**p* = 0.0168; tdTomato, *p* = 0.9878; ChR2, \*\**p* = 0.0045; (M) PF→CPuTRAPed, F_(1, 16)_ = 0.06721, p = 0.7987; tdTomato, p = 0.8915; ChR2, p = 0.6813. tdTomato, *N* = 9; ChR2, *N* = 9. Two-way ANOVA followed by Sidak’s post hoc test. (E, N) Stimulation paradigm (alternating 60-s OFF/ON epochs) and representative snapshots from illustrating behavioral phenotype evoked by M2→CPu^TRAPed^ (E) or PF→CPu^TRAPed^ (N) activation. (F–H, O–Q) AIM subtype scores (axial, limb, and orolingual) during OFF versus ON epochs for M2→CPu^TRAPed^ (F–H) or PF→CPu^TRAPed^ (O–Q) stimulation. (F, O) Axial AIM score. (F) M2→CPu^TRAPed^, F_(1, 16)_ = 1.802, *p* = 0.1982; tdTomato, *p* = 0.9368; ChR2, *p* = 0.08. (O) PF→CPu^TRAPed^, F_(1, 16)_ = 0.7800, *p* = 0.3902; tdTomato, *p* = 0.8478; ChR2, *p* = 0.1826. (G, P) Limb AIM score. (G) M2→CPu^TRAPed^, F_(1, 16)_ = 5.298, ^*^*p* = 0.0351; tdTomato, *p* > 0.9999; ChR2, \*\**p* = 0.0099. (P) PF→CPu^TRAPed^, F_(1, 16)_ = 0.6154, *p* = 0.4442; tdTomato, *p* = 0.2178; ChR2, *p* = 0.8292. (H, Q) Orolingual AIM score. (H) M2→CPu^TRAPed^, F_(1, 16)_ = 8.745, \*\**p* = 0.0093; tdTomato, *p* = 0.8164; ChR2, \*\**p* = 0.0047. (Q) PF→CPu^TRAPed^, F_(1, 16)_ = 0.001278, *p* = 0.9719; tdTomato, *p* = 0.7076; ChR2, *p* = 0.6756. tdTomato, *N* = 9; ChR2, *N* = 9. Two-way ANOVA followed by Sidak’s post hoc test. (I, R) Global AIM score during M2→CPu^TRAPed^ (I) stimulation with low-dose levodopa or PF→CPu^TRAPed^ (R) stimulation without levodopa. (I) M2→CPu^TRAPed^, F_(1, 11)_ = 9.766, \*\**p* = 0.0097; tdTomato, *p* = 0.9193; ChR2, \*\*\**p* = 0.0004. tdTomato, *N* = 5; ChR2, *N* = 8. (R) PF→CPu^TRAPed^, F_(1, 16)_ = 0.2751, *p* = 0.6071; tdTomato, *p* = 0.2441; ChR2, *p* = 0.6482. tdTomato, *N* = 9; ChR2, *N* = 9. Two-way ANOVA followed by Sidak’s post hoc test. *N*, animals. Data are mean ± SEM.

Consistent with the anatomical alignment observed in M2, stimulation of ensemble-projecting M2 neurons had little effect on open-field locomotion but was sufficient to evoke dyskinesia-like AIMs in the absence of levodopa (Figs. 6A–F and S10C). This effect was most prominent in limb and orolingual AIMs (Fig. 6G–I), indicating that the dyskinesia-promoting influence of M2 is carried, at least in part, by a direct M2→ensemble channel. When dyskinesia was primed with a low dose of levodopa, activation of the same M2→ensemble subcircuit further amplified AIMs across subtypes (Figs. 6I and S10D–F), suggesting that levodopa-dependent sensitization lowers the threshold for cortical recruitment of the ensemble.

In contrast, selective stimulation of PF neurons that directly innervate the striatal ensemble failed to modify behavior. PF→ensemble activation did not reproduce the locomotor suppression seen with projection-wide PF→CPu stimulation, nor did it alleviate levodopa-evoked dyskinesia (Figs. 6L–Q and S10H). PF→ensemble activation was also ineffective in the absence of levodopa (Figs. 6R and S10I–K). This broad-versus-restricted dissociation aligns with the topographic dissociation identified above: PF neurons projecting to the ensemble are spatially segregated from the PF population preferentially engaged during dyskinesia. Accordingly, the anti-dyskinetic impact of projection-wide PF activation is unlikely to be mediated by direct PF→ensemble recruitment; instead, it points to an ensemble-external thalamostriatal brake, potentially implemented through PF inputs to non-ensemble striatal targets and/or local inhibitory microcircuits. These circuit-restricted perturbations therefore dissociate an ensemble-aligned M2 driver from a topographically and functionally distinct PF brake.

### Ensemble-restricted *Grin1* knockdown reduces peak dyskinesia and M2-driven dyskinesia

Because M2→ensemble synapses exhibited a prominent NMDAR component and because selective M2→ensemble activation promoted dyskinesia, we asked whether NMDAR signaling within dyskinesia-linked ensemble neurons is required for efficient cortical recruitment. We delivered a Cre-dependent shRNA targeting *Grin1*, which encodes the obligate GluN1 subunit of the NMDAR, or a scramble control to the dorsal striatum and used FosTRAP-dependent recombination during LID to initiate ensemble-restricted knockdown (Figs. 7A, B and S11A). Before recombination (“pre-knockdown”), scramble and *Grin1*-shRNA cohorts were indistinguishable in spontaneous locomotion and levodopa-evoked dyskinesia severity (Fig. S11B–H). Likewise, M2 terminal stimulation evoked comparably robust dyskinesia-like AIMs in both groups before knockdown induction (Fig. S11I–M), establishing matched baseline responsiveness to M2 drive.

**Fig. 7:**
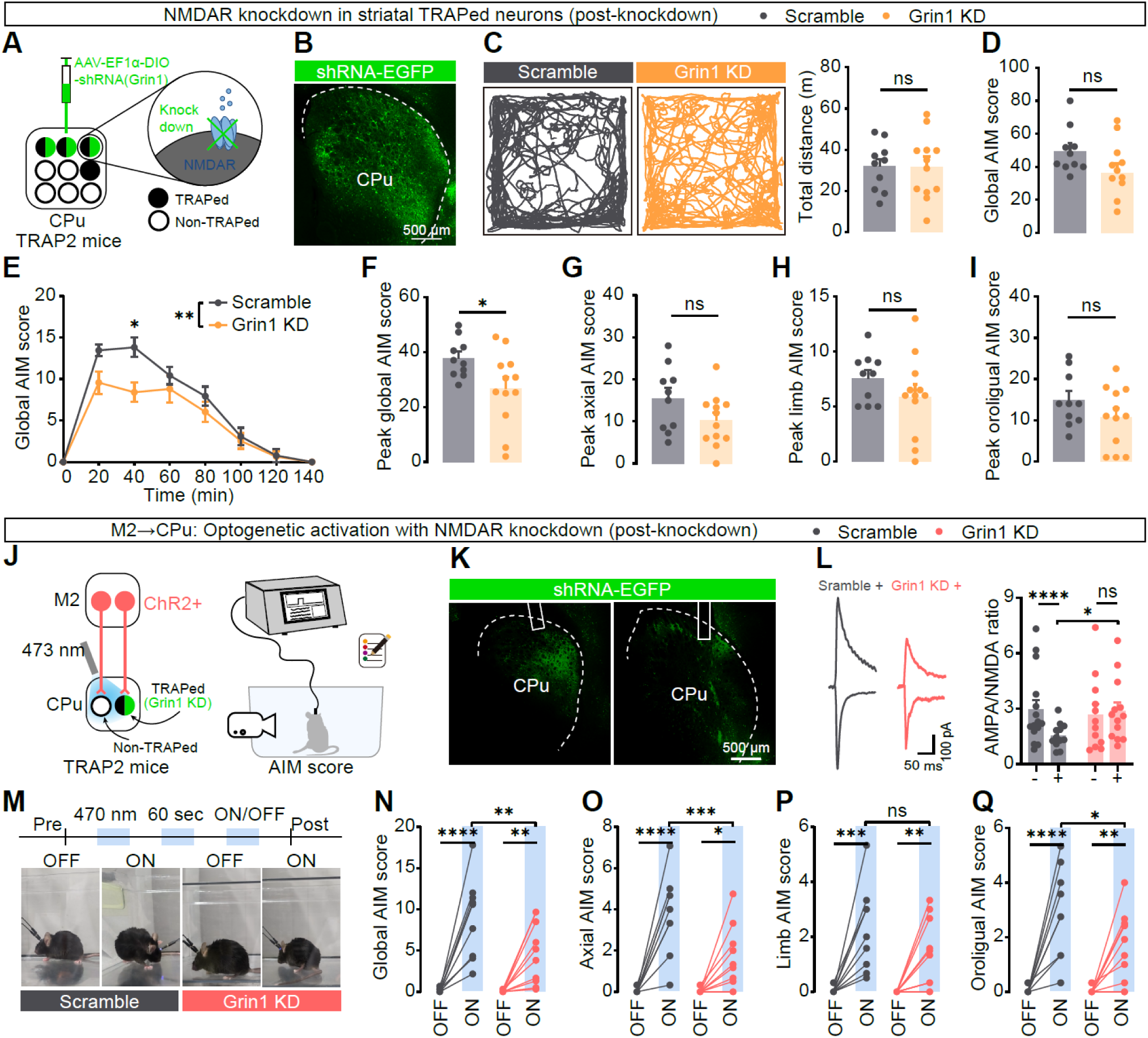
Activity-dependent NMDAR knockdown in dyskinesia-linked striatal ensembles reduces peak dyskinesia and weakens M2-driven dyskinesia. (A) Strategy for ensemble-restricted NMDAR knockdown. A Cre-dependent AAV expressing *Grin1* shRNA–EGFP (*Grin1* KD) or a matched scramble shRNA control was injected into the dorsal striatum (CPu) of TRAP2 mice. Following LID-associated TRAP induction (4-OHT; “post-knockdown”), shRNA expression—and thus GluN1/NMDAR reduction—was confined to FosTRAP-tagged ensemble neurons. (B) Representative coronal section showing *Grin1* shRNA–EGFP expression in CPu after TRAP induction. (C) Representative 20-min open filed trajectories (left) and total distance traveled (right) in Scramble and *Grin1* KD mice. Scramble, *N* = 10; *Grin1* KD, *N* = 12. ns, *p* = 0.9821; two-tailed unpaired Student’s *t* test. (D) Cumulative dyskinesia severity across the levodopa scoring session, expressed as the global AIM score summed across monitoring epochs. Scramble, *N* = 10; *Grin1* KD, *N* = 12. ns, *p* = 0.0795; two-tailed unpaired Student’s *t* test. (E) Time courses of global AIM score following levodopa administration, showing a reduction that is most apparent during the peak dyskinesia window. F_(7, 140)_ = 3.147, \*\**p* = 0.0032; 40 min, \**p* = 0.0309. Scramble, *N* = 10; *Grin1* KD, *N* = 12. \**p* = 0.0309, Two-way ANOVA followed by Sidak’s post hoc test. (F–I) Peak AIM scores for global AIM (F) and axial (G), limb (H), and orolingual (I) sub-score. (F) Global, \**p* = 0.0318. (G) Axial, *p* = 0.102. (H) Limb, *p* = 0.2142. (I) Orolingual, *p* = 0.1801. Scramble, *N* = 10; *Grin1* KD, *N* = 12. Two-tailed unpaired Student’s *t* test. (J) Design for testing whether ensemble NMDAR knockdown reduces dyskinesia driven by projection-wide M2 terminal activation in striatum. ChR2 was expressed in M2, and terminals were stimulated in CPu while AIMs were scored. (K) Representative images verifying striatal shRNA expression and optic-fiber placement for M2 terminal stimulation. (L) Electrophysiological validation of knockdown in TRAPed neurons, quantified as an increased AMPA/NMDA ratio at M2→TRAPed synapses after *Grin1* KD. Scramble−, *n* = 15, *N* = 5; scramble+, *n* = 12, *N* = 5; *Grin1* KD−, *n* = 12, *N* = 4; *Grin1* KD+, *n* = 13, *N* = 4. F_(1, 22)_ = 13.22, \*\**p* = 0.0015; Scramble− vs. Scramble+, \*\*\*\**p* < 0.0001; *Grin1* KD− vs. *Grin1* KD+, ns, *p* = 0.4538; Scramble+ vs. *Grin1* KD+, \**p* = 0.0246. Two-way ANOVA followed by Sidak’s post hoc test. (M) Stimulation paradigm (60-s OFF/ON epochs) and representative snapshots illustrating behavior during M2 terminal stimulation in Scramble and *Grin1* KD mice. (N–Q) Global (N) and subtype (O–Q) AIM scores during paired OFF and ON epochs of M2 terminal stimulation. (N) Global, F_(1, 17)_ = 4.689, \**p* = 0.0449; Scramble, *N* = 9 mice, \*\*\*\**p* < 0.0001; *Grin1* KD, *N* = 10 mice, \*\**p* = 0.007. Scramble_ON vs. *Grin1* KD_ON, \*\**p* = 0.0082. (O) Axial, F_(1, 17)_ = 7.617, \**p* = 0.0134; Scramble, \*\*\*\**p* < 0.0001; *Grin1* KD, \**p* = 0.0202; Scramble_ON vs. *Grin1* KD_ON, \*\*\**p* = 0.001. (P) Limb, F_(1, 17)_ = 1.558, *p* = 0.2289; Scramble, \*\*\**p* = 0.0003; *Grin1* KD, \*\**p* = 0.0084; Scramble_ON vs. *Grin1* KD_ON, *p* = 0.1453. (Q) Orolingual, F_(1, 17)_ = 3.334, *p* = 0.0855; Scramble, \*\*\*\**p* < 0.0001; *Grin1* KD, \*\**p* = 0.009; Scramble_ON vs. *Grin1* KD_ON, \**p* = 0.0248. Scramble, *N* = 9; *Grin1* KD, *N* = 10. Two-way ANOVA followed by Sidak’s post hoc test. *n,* cells*. N*, animals. Data are as mean ± SEM.

After recombination (“post-knockdown”), ensemble-restricted *Grin1* knockdown did not measurably alter spontaneous locomotion in the open field (Fig. 7C), and the overall session-level global AIM score showed only a modest, non-significant reduction (Fig. 7D). However, time-course analysis revealed that the effect of *Grin1* knockdown was concentrated in the peak dyskinesia window after levodopa administration (Fig. 7E). Consistent with this peak-weighted effect, *Grin1* knockdown significantly reduced the peak global AIM score (Fig. 7F), whereas individual subtype peak scores trended downward without reaching significance when analyzed separately (Fig. 7G–I). Thus, reducing NMDAR availability selectively within dyskinesia-linked ensemble neurons preferentially dampened peak LID expression.

We next tested whether ensemble NMDAR signaling contributes to M2-driven dyskinesia. After initiating *Grin1* knockdown, we optogenetically activated M2 terminals in dorsal striatum (Fig. 7J and K) and confirmed reduced NMDAR contribution at corticostriatal synapses onto TRAPed neurons by post hoc electrophysiology, reflected in an increased AMPA/NMDA ratio (Fig. 7L). Under these conditions, M2 terminal stimulation still evoked dyskinesia-like behaviors in both cohorts, but *Grin1* knockdown significantly attenuated the induced AIMs. Compared with scramble controls, *Grin1* knockdown mice showed a reduced global AIM response to M2 stimulation (Fig. 7M and N), with significant attenuation of axial and orolingual components (Fig. 7O and Q), whereas the limb component was less affected (Fig. 7P). Together with the matched pre-knockdown baseline, these findings identify NMDAR signaling within dyskinesia-linked striatal ensembles as an important postsynaptic substrate enabling M2 inputs to drive dyskinesia expression.

## Discussion

Our study identifies two opposing circuit mechanisms that control LID: a motor cortical drive that recruits dyskinesia-linked striatal ensemble neurons and a parafascicular thalamostriatal brake that suppresses ongoing dyskinesia. Previous work established that only a subset of striatal projection neurons is causally engaged during LID,^16–19^ but the upstream pathways that recruit or restrain this pathological population remained unclear. Here, M2 and PF emerged as dominant afferents with opposite behavioral effects. Projection-wide activation of M2 promoted or exacerbated axial, limb, and orolingual abnormal involuntary movements, whereas projection-wide activation of PF suppressed ongoing dyskinesia and shifted animals toward non-dyskinetic behavioral states. Chronic levodopa preserved effective pathway-to-ensemble coupling despite an overall loss of presynaptic terminals, but imposed distinct synaptic mechanisms: M2 inputs became biased toward NMDAR-weighted excitation and spatial dispersion, whereas PF influence was associated with stronger polysynaptic inhibition. Together, these findings support a preclinical circuit framework in which LID reflects an imbalance between an ensemble-aligned cortical driver and an ensemble-external thalamostriatal brake (Fig. 8).

**Fig. 8:**
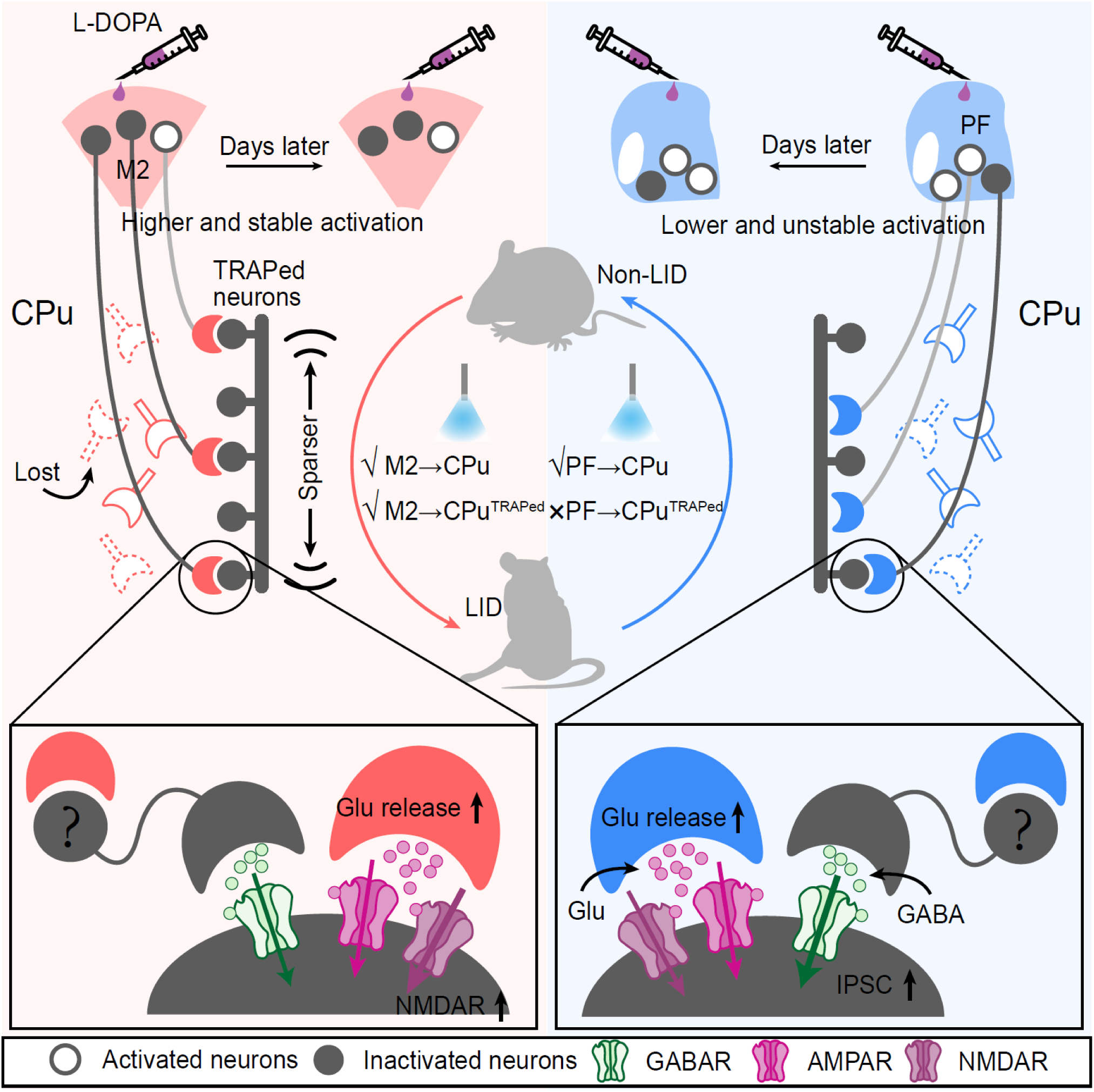
Working model of opponent cortical and thalamic access to dyskinesia-linked striatal ensembles in levodopa-induced dyskinesia. Chronic levodopa biases access to dyskinesia-linked ensemble neurons in the dorsal striatum (CPu) through two dominant long-range pathways with opposite functional roles. Left, M2: dyskinesia preferentially re-engages the subset of secondary motor cortex (M2) neurons that project to the striatal ensemble, producing relatively stable cortical access to ensemble neurons across dyskinesia episodes. Although overall M2 terminal availability is reduced, effective M2-to-ensemble coupling is preserved, contacts become more spatially dispersed, and synaptic integration becomes biased toward NMDAR-weighted excitation, consistent with enhanced cortical recruitment of the ensemble. Accordingly, both projection-wide M2→CPu stimulation and circuit-restricted M2→CPu^TRAPed^ stimulation promote dyskinesia. Right, PF: parafascicular thalamic (PF) input exerts the opposite behavioral effect. Broad PF→CPu activation suppresses dyskinesia and favors a non-dyskinetic state, but PF neurons directly innervating the ensemble are not preferentially reactivated during dyskinesia, indicating an activity–connectivity dissociation. At the synaptic level, PF contacts onto ensemble neurons remain relatively stable in spatial organization but recruit stronger polysynaptic inhibition, shifting net synaptic drive toward inhibition. In contrast to M2, circuit-restricted PF→CPu^TRAPed^ activation is ineffective, indicating that PF suppression is implemented largely through an ensemble-external thalamostriatal brake. Lower insets summarize the dominant synaptic rules inferred for each pathway. Question marks denote unresolved intermediary circuit elements and/or postsynaptic cell-type specificity. Dashed boutons indicate terminals lost after chronic levodopa exposure.

A recurring structural theme across our data is that chronic levodopa appears to preserve access to dyskinesia-linked ensemble neurons even as broader presynaptic terminal fields contract. Both M2 and PF showed reduced or trending reductions in presynaptic bouton abundance within striatum, yet the number and density of putative contacts onto ensemble dendrites were maintained. This pattern is most consistent with preferential maintenance and/or biased reallocation of surviving cortical and thalamic contacts onto ensemble targets, potentially at the expense of inputs to non-ensemble neurons. Such ensemble-biased access provides a circuit-level explanation for how repeated levodopa exposure might stabilize a pathological motor state without globally increasing excitatory innervation throughout the striatum.

This finding raises a broader question: why would levodopa exposure in a dopamine-denervated striatum simultaneously erode overall long-range innervation, preserve effective coupling onto a dyskinesia-linked ensemble, and reorganize the spatial arrangement of those inputs? A plausible explanation is that chronic levodopa creates a non-physiological reinforcement regime for striatal plasticity. Because levodopa has a short half-life and dopamine buffering is diminished after denervation, receptor activation becomes abnormally pulsatile, a temporal pattern closely linked to molecular and structural adaptations associated with dyskinesia.^5,13^ Under such pulsatile drive, repeated dopamine peaks may repeatedly select the same striatal ensemble and stabilize the afferent inputs that successfully recruit it, thereby preserving privileged access to the ensemble even as the broader afferent landscape contracts. This interpretation is consistent with the clinical rationale for continuous dopaminergic stimulation, which aims to reduce motor complications by smoothing receptor stimulation and limiting pulse-driven plasticity.^5^

Motor learning provides a useful counterpoint for interpreting the spatial organization of ensemble inputs. During physiological learning, cortical inputs to striatal engram neurons are selectively strengthened and can become spatially clustered along dendrites, a configuration thought to promote efficient nonlinear integration of aligned drive.^29,37,40,41^ Recent *in vivo* imaging further shows that motor learning remodels corticostriatal boutons and increases spatial clustering, whereas thalamostriatal boutons are comparatively stable.^42^ Against this adaptive template, the increased dispersion of M2-to-ensemble contacts that we observe after chronic levodopa—despite preserved effective coupling—may represent a homeostatic attempt to dilute cooperative cortical excitation when ensemble drive becomes excessive. In this view, LID may reflect maladaptive consolidation of ensemble recruitment under pulsatile dopamine reinforcement, with synaptic dispersion serving as an insufficient compensatory response rather than a learning-like optimization.

Functionally, M2 and PF shared one broad feature but diverged at the point most relevant to behavior. Both M2→ensemble and PF→ensemble pathways showed altered paired-pulse dynamics, consistent with enhanced effective transmission during repeated activation. This convergence is notable because cortical and thalamic glutamatergic inputs are classically distinguished by different short-term dynamics and presynaptic operating regimes.^43^ Thus, chronic levodopa may partially override canonical pathway differences at the level of ensemble access. However, because these short-term dynamics were measured using optogenetically evoked responses, they should be interpreted as evidence of altered effective transmission rather than a definitive measure of presynaptic release probability. The fact that M2 and PF produced opposite behavioral outcomes despite partially convergent short-term dynamics indicates that the critical divergence lies in how ensemble neurons decode these inputs postsynaptically and how each pathway engages local striatal circuitry.

For M2 inputs, chronic levodopa altered the quality of excitatory integration rather than producing a large change in evoked excitation-to-inhibition balance. M2→ensemble contacts became disproportionately NMDAR-weighted, reflected by a lower AMPA/NMDA ratio relative to neighboring non-ensemble neurons. Within classical corticostriatal frameworks, NMDAR-rich integration is well suited to support depolarized striatal state transitions and sustained ensemble engagement.^43^ Our ensemble-restricted *Grin1* knockdown provides causal support for the relevance of this mechanism: reducing NMDAR availability selectively in TRAPed ensemble neurons left baseline locomotion largely intact, but reduced peak dyskinesia and weakened the dyskinesia-promoting effect of M2 input activation. These findings do not identify NMDARs as a single molecular switch for LID. Instead, they support the idea that NMDAR-weighted corticostriatal integration acts as a permissive gain axis that enables M2 input to recruit the dyskinesia-linked ensemble more effectively.

This circuit-level axis is compatible with, but should not be equated with, the clinical efficacy of glutamatergic strategies for dyskinesia. Amantadine is the best-established example and is often discussed in relation to NMDAR antagonism.^44^ However, its anti-dyskinetic actions are unlikely to be explained solely by NMDAR blockade, and alternative postsynaptic mechanisms, including Kir2 channel-linked effects, have been proposed.^45^ Preclinical work also supports broader glutamate receptor modulation, including AMPAR antagonism and combined AMPA/NMDA blockade, in reducing LID or related motor complications.^46–49^ Our data therefore identify NMDAR-weighted cortical recruitment of dyskinesia-linked ensemble neurons as one modifiable synaptic axis, while leaving open the possibility that additional glutamatergic and non-glutamatergic mechanisms shape the full dyskinesia phenotype.

PF inputs followed a different functional logic. Although intralaminar thalamostriatal projections are glutamatergic and often associated with salience or arousal signaling,^35,38^ PF activation in our recordings recruited disproportionately strong feedforward inhibition, shifting net synaptic drive toward inhibition. This inhibitory routing may help explain how a glutamatergic thalamic input can suppress dyskinesia. PF may engage local inhibitory motifs that sharpen action selection, stabilize striatal competition, and limit runaway recruitment of dyskinesia-generating ensemble activity. Whether this PF-linked inhibition is maladaptive—contributing to abnormal gating—or compensatory—serving to restrain cortical overdrive—remains unresolved. Clinical observations that benzodiazepines can lessen levodopa-induced diphasic dyskinesias and that midazolam may transiently suppress severe choreiform dyskinesia, although confounded by sedation, are conceptually consistent with an inhibitory brake mechanism.^50,51^ Together with the M2 NMDAR-weighted recruitment axis, these findings suggest that chronic levodopa installs pathway-specific integration regimes: cortical inputs become biased toward ensemble recruitment, whereas PF influence is preferentially routed through inhibition.

A particularly important dissociation emerged from the PF experiments. Projection-wide PF→striatum activation reduced AIM scores and, in 3D behavioral analyses, shifted animals away from dyskinetic motifs toward a more non-dyskinetic repertoire. This was not simply a reduction in movement; PF stimulation reduced occupancy of severe dyskinetic states while allowing re-expression of locomotor behaviors. By contrast, when PF input was restricted to ensemble-defined monosynaptic connectivity using rabies-dependent targeting, selective PF→ensemble activation failed to modify dyskinesia. This contrast indicates that PF does not suppress dyskinesia by directly engaging the same ensemble-targeting channel that it anatomically contributes to.

Our anatomical data provide a parsimonious explanation for this broad-versus-restricted dissociation. In PF, neurons most strongly reactivated during dyskinesia were spatially segregated from PF neurons that project monosynaptically to the dyskinesia-linked striatal ensemble. Thus, the PF module carrying anti-dyskinetic leverage appears to be different from the PF module that directly innervates the ensemble. More broadly, this suggests a disease-relevant principle for basal ganglia organization: although cortico–basal ganglia–thalamic circuits are often described as parallel and topographically matched,^21,22^ pathological functional maps can diverge from canonical anatomical projection maps. For circuit-targeted therapies, this point is important. Anatomical connectivity alone may not be sufficient to identify the circuit module that controls a pathological state; disease-linked activity and behavioral effect must also be considered.

These results also have potential implications for neuromodulation-based strategies. The opposing effects of M2 and PF suggest that LID may be modifiable by targeting either excessive cortical recruitment or insufficient thalamostriatal restraint. Clinical studies have reported that transcranial direct current stimulation over motor cortex can reduce LID symptoms in some patients,^52^ although whether M2 or related premotor cortical territories represent optimal targets remains unknown. Deep brain stimulation of the centromedian/parafascicular complex has shown anti-dyskinetic effects in some PD contexts while improving tremor,^53^ and chronic PF stimulation reduces dyskinesia in rodent models.^54^ Our data provide a mechanistic rationale for considering PF-related thalamostriatal circuits as candidate anti-dyskinetic targets. However, the dissociation between projection-wide PF suppression and selective PF→ensemble ineffectiveness also cautions that stimulation site, recruited PF submodule, and downstream striatal microcircuit engagement are likely to be critical determinants of therapeutic efficacy.

In summary, this study advances the dyskinesia-ensemble framework by identifying upstream circuit mechanisms that recruit and restrain pathological striatal ensemble activity. Chronic levodopa promotes ensemble-biased preservation of pathway access and imposes distinct synaptic mechanisms on two dominant afferents: M2 provides an NMDAR-weighted cortical drive capable of aggravating dyskinesia through ensemble-targeting subcircuits, whereas PF provides an ensemble-external thalamostriatal brake likely implemented through inhibitory routing and modular thalamostriatal organization. These findings define LID as a targetable circuit imbalance between cortical recruitment and thalamostriatal restraint, with implications for future neuromodulation and synapse-informed therapeutic strategies.

### Limitations of the study

Several limitations should be considered. *First*, although our FosTRAP strategy provides activity-dependent access to a dyskinesia-linked striatal population, it does not by itself resolve the full cell-type composition of that ensemble. Prior work indicates enrichment for direct-pathway neurons in dyskinesia-linked TRAPed populations,^17^ but the relative contributions of direct-pathway MSNs, indirect-pathway MSNs, and striatal interneurons in the present experiments remain to be defined. This is particularly important for interpreting PF-linked inhibitory routing, because thalamostriatal inputs can engage multiple striatal cell types, including direct- and indirect-pathway projection neurons, cholinergic interneurons, fast-spiking interneurons, and other GABAergic interneuron classes. Future work combining cell-type-resolved tagging, recording, and perturbation will be required to assign the cortical drive and PF brake to specific striatal cell populations.

*Second*, the intermediary microcircuits that convert PF glutamatergic input into a net inhibitory influence on ensemble neurons remain unresolved. The striatum contains multiple inhibitory substrates capable of implementing feedforward inhibition, including fast-spiking interneurons, other GABAergic interneurons, and recurrent MSN collateral interactions. These microcircuits are diverse, dynamically regulated by dopamine state, and increasingly implicated in movement disorders.^55–63^ Dopamine can modulate local GABAergic inhibition onto striatal MSNs,^64–66^ providing one possible bridge between levodopa exposure and strengthened PF-linked inhibition. In addition, thalamostriatal synapses are target-specific and state-dependent:^67–74^ thalamostriatal remodeling can bias striatal drive in parkinsonian mice with consequences for motor impairment,^75,76^ and cholinergic signaling can selectively amplify PF-driven excitation of iMSNs.^77^ These observations raise the possibility that PF’s anti-dyskinetic effect depends on differential regulation of direct- and indirect-pathway output through local microcircuits rather than a simple PF→ensemble monosynaptic mechanism. Determining whether the enhanced inhibitory routing observed here is part of the pathological state or a compensatory protection against cortical overdrive will require causal dissection of these intermediary circuit elements.

*Third*, although ensemble-restricted *Grin1* knockdown supports a role for NMDAR-dependent cortical recruitment, the specific NMDAR subunit composition and downstream signaling mechanisms remain unknown. NMDAR properties depend strongly on subunit assembly, which shapes synaptic integration, plasticity rules, and pharmacological sensitivity.^78,79^ Altered NMDAR subunit composition could therefore be a key determinant of cortical input gain during LID and may influence which glutamatergic interventions are most effective. *Finally*, our study was performed in a unilateral 6-OHDA mouse model with viral and optogenetic tools. This approach provides causal circuit resolution but does not fully capture the progressive, multisystem nature of human PD or the clinical heterogeneity of LID. Validation in additional models and, where possible, in human-relevant systems will be important for translation.

## Contributors

J.W., X.-Y.T., Z.L., W.-G.L., and L.S. conceived the project, designed the experiments, and interpreted the results. J.W. and X.-Y.T. performed the most experiments including behavioral experiments, animal surgery, immunofluorescence, populational Ca^2+^ and patch clamp recording and data analysis. J.-X.L. and J.-Y.S. commented on data interpretation. W.X., Y.-L.W., X.Y., and Y.-J.W. provided suggestions and guidance for some experimental operations. X.-N.L. commented on the manuscript. Y.-X.L. performed genotyping. J.W., X.-Y.T., Z.L., W.-G.L., and L.S. wrote the manuscript with contributions from all authors. All authors read and approved the final manuscript.

## Data sharing statement

Any additional information required to reanalyze the data reported in this paper is available from the lead contact upon reasonable request. Proposals should be directed to (L.S.).

## Acknowledgements

This study was supported by grants from the STI2030-Major Projects (2022ZD0208605), the National Natural Science Foundation of China (82171242, 82471260, 82271274, and 82474606). During the preparation of this work, the author(s), as the non-native English speakers, used ChatGPT 5.4 (OpenAI) to improve the language and enhance its readability. The author(s) have reviewed and confirmed the validity of the text and take(s) full responsibility for the content of the publication.

## Appendix A. Supplementary data

Supplementary data including 11 Supplemental Figures related to this article can be found online.

## Supplemental Figures and Figure Legends

**Fig. S1:**
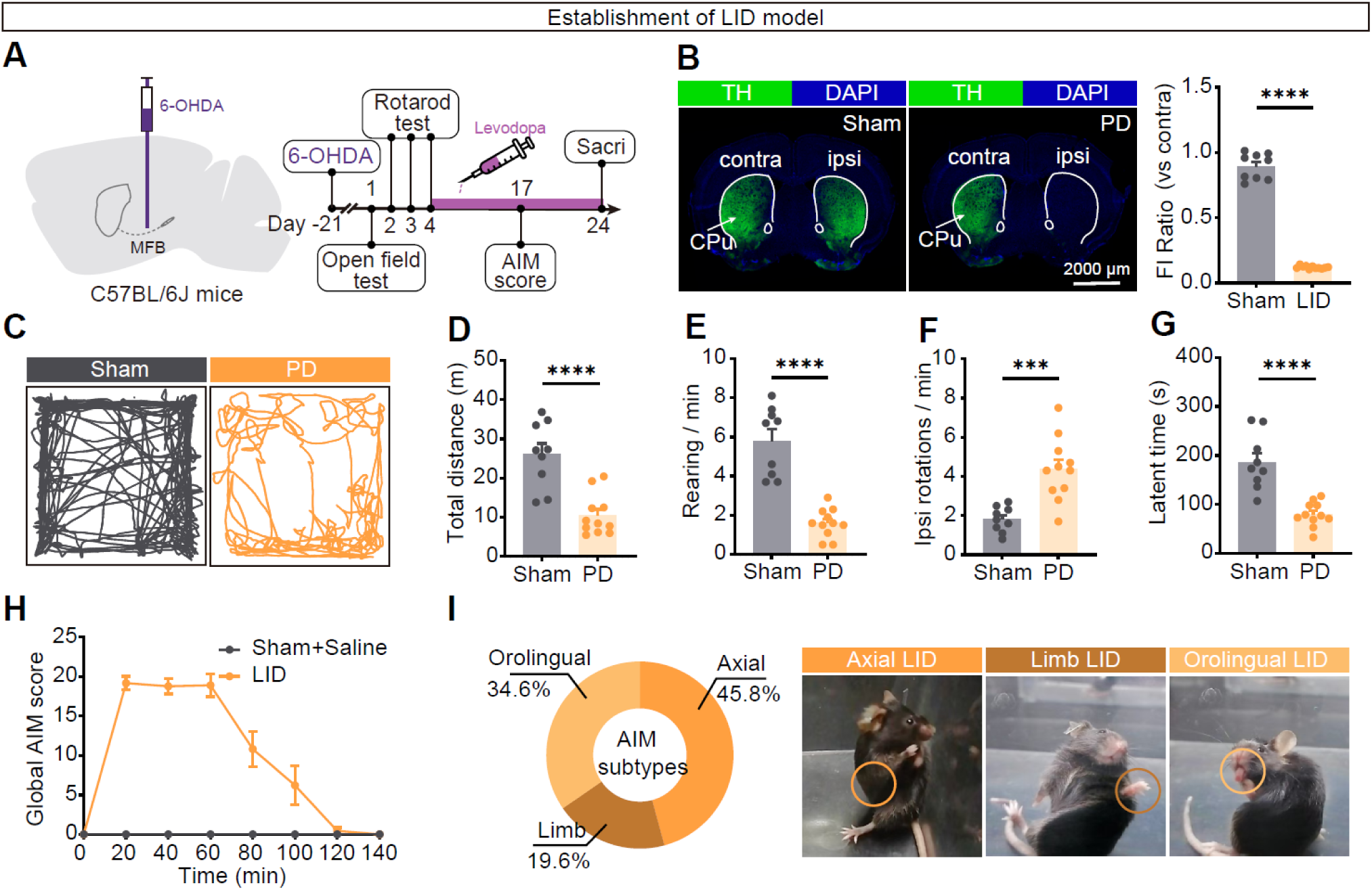
Unilateral 6-OHDA lesioning and levodopa priming establish a robust dyskinesia model. (A) Experimental design and timeline. Mice received a unilateral 6-OHDA injection into the medial forebrain bundle (MFB) to induce dopamine depletion. Parkinsonian motor deficits were assessed by open field and accelerating rotarod, followed by chronic levodopa priming and abnormal involuntary movement (AIM) scoring. (B) Left, representative coronal sections of striatum showing tyrosine hydroxylase (TH) immunoreactivity in sham and 6-OHDA–lesioned mice. Dopaminergic innervation is markedly reduced in the ipsilateral (ipsi) striatum relative to the contralateral (contra) hemisphere in lesioned mice. Right, quantification of the TH fluorescence intensity ratio (ipsi/contra). Sham, *N* = 9; PD, *N* = 11. \*\*\*\**p* < 0.0001; two-tailed unpaired Student’s *t* test. (C) Representative center-point trajectories during a 10-min open field session in sham (left) and6-OHDA–lesioned (right) mice. (D– F) Open field quantification showing reduced locomotor distance (D), reduced rearing frequency (E), and increased ipsilateral rotations (F) in 6-OHDA–lesioned mice relative to sham controls. (D) \*\*\**p* = 0.001, Mann-Whitney U test. (E) \*\*\*\**p* < 0.0001, two-tailed unpaired Student’s *t* test. (F) \*\*\**p* = 0.0003; two-tailed unpaired Student’s *t* test. Sham, *N* = 9; PD, *N* = 11. (G) Accelerating rotarod performance (latency to fall) demonstrating impaired motor coordination in 6-OHDA–lesioned mice. Sham, *N* = 9; PD, *N* = 11. \*\*\*\**p* < 0.0001, two-tailed unpaired Student’s *t* test. (H) Time course of global (AIM) score after levodopa administration in PD mice compared with saline-injected sham controls, demonstrating robust LID. (I) Left, proportion of axial, limb, and orolingual components contributing to the global AIM score. Right, representative snapshots illustrating the three AIM subtypes (axial: trunk/neck posturing and twisting; limb: contralateral forelimb movements; orolingual: jaw/tongue/facial movements; orange circles indicate the affected body region). *N*, animals. Data are mean ± SEM.

**Fig. S2:**
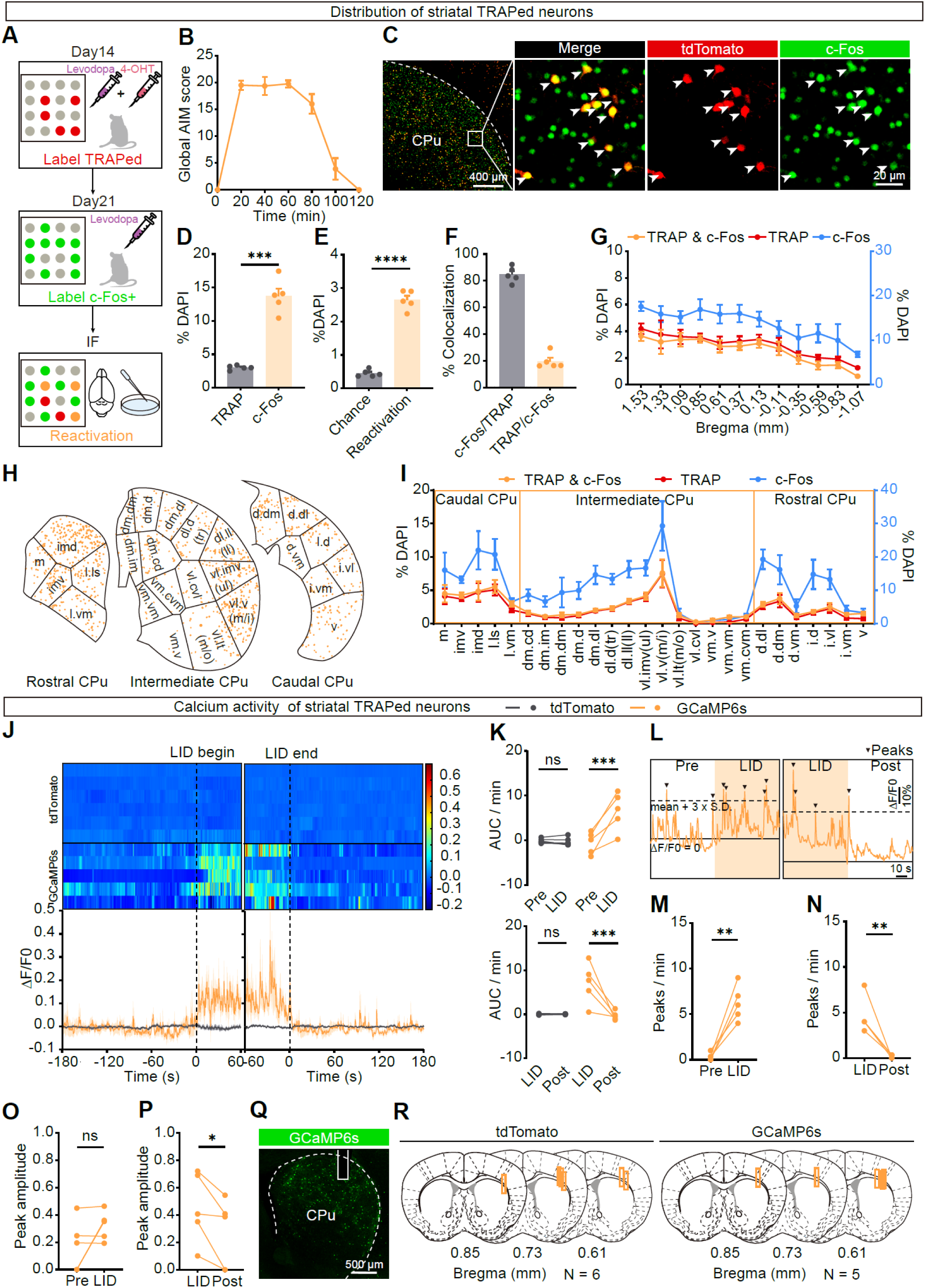
FosTRAP identifies a reproducibly reactivated striatal ensemble with dyskinesia-locked Ca^2+^ dynamics. (A) Experimental design for activity-tagging and reactivation analysis in TRAP2::Ai14 mice. During the first L-DOPA–evoked dyskinesia episode, 4-hydroxytamoxifen (4-OHT) was administered to permanently label active neurons as tdTomato^+^ (TRAPed). One week later, mice received a second L-DOPA challenge and were perfused for c-Fos immunostaining; tdTomato^+^c-Fos^+^ cells were scored as reactivated ensemble neurons. (B) Time course of global AIM score during the levodopa session before TRAP induction. (C) Representative images of dorsolateral CPu showing TRAPed neurons (tdTomato, red), LID-reactivated neurons (c-Fos, green), and overlapped neurons (merge). Arrowheads denote reactivated ensemble neurons (tdTomato^+^c-Fos^+^). (D) Density of TRAPed neurons and c-Fos^+^ neurons in CPu, expressed as the percentage of DAPI^+^ nuclei. *N* = 5. \*\*\**p* = 0.0006; two-tailed paired Student’s *t* test. (E) Reactivation enrichment compared with chance overlap. *N* = 5. \*\*\*\**p* < 0.0001; two-tailed paired Student’s *t* test. (F) Reactivation expressed as (i) the fraction of TRAPed neurons that were c-Fos^+^ (c-Fos^+^/TRAP^+^) and (ii) the fraction of c-Fos^+^ neurons captured by FosTRAP (TRAP^+^/c-Fos^+^). *N* = 5. (G) Rostrocaudal distribution of striatal TRAPed neurons, c-Fos^+^ neurons, and overlapped neurons (densities as %DAPI; Bregma coordinates indicated). *N* = 5. (H) Representative reconstructions of TRAPed neuron distributions across striatal functional domains at rostral, intermediate, and caudal CPu levels (orange spots). (I) Domain-level quantification of TRAPed neurons, c-Fos^+^ neurons, and overlapped neurons across medial–lateral striatal territories, plotted separately for caudal, intermediate, and rostral CPu (densities as %DAPI). *N* = 5. (J) Fiber photometry of ensemble activity in TRAP2::Ai162 mice expressing GCaMP6s. Heatmaps (rows, individual mice) and mean traces (bottom) show ΔF/F aligned to dyskinesia onset (“LID begin”) and offset (“LID end”). tdTomato recordings from TRAP2::Ai14 mice are shown as a motion/bleaching control (gray). *N* = 5. (K) Quantification of ΔF/F area under the curve (AUC) per min during LID emergence (top) and during LID decay. Top, F_(1, 9)_ = 28.14, \*\*\**p* = 0.0005; tdTomato, *p* = 0.9767; GCaMP6s, \*\*\**p* = 0.0001. Bottom, F_(1, 9)_ = 15.87, \*\**p* = 0.0032; tdTomato, *p* = 0.999; GCaMP6s, \*\*\**p* = 0.0009. tdTomato, *N* = 6; GCaMP6s, *N* = 5. Two-way ANOVA followed by Sidak’s post hoc test. (L) Representative ΔF/F trace illustrating calcium event detection; peaks (triangles) were defined as excursions exceeding mean (ΔF/F) + 3 SD. *N* = 5. (M, N) Peak frequency increased during LID relative to pre-LID (M, \*\**p* = 0.0033) and decreased after LID offset (N, \*\**p* = 0.009, two-tailed paired Student’s *t* test). *N* = 5. (O, P) Peak amplitude was not significantly altered at LID onset (O, *p* = 0.1727) but decreased after LID offset (P, \**p* = 0.0355). *N* = 5. Two-tailed paired Student’s *t* test. (Q) Representative section confirming GCaMP6s expression and optical fiber placement in CPu. (R) Summary of photometry fiber placements across mice (orange rectangles) overlaid on atlas sections. *N* = 5. *N*, animals. Data are mean ± SEM.

**Fig. S3:**
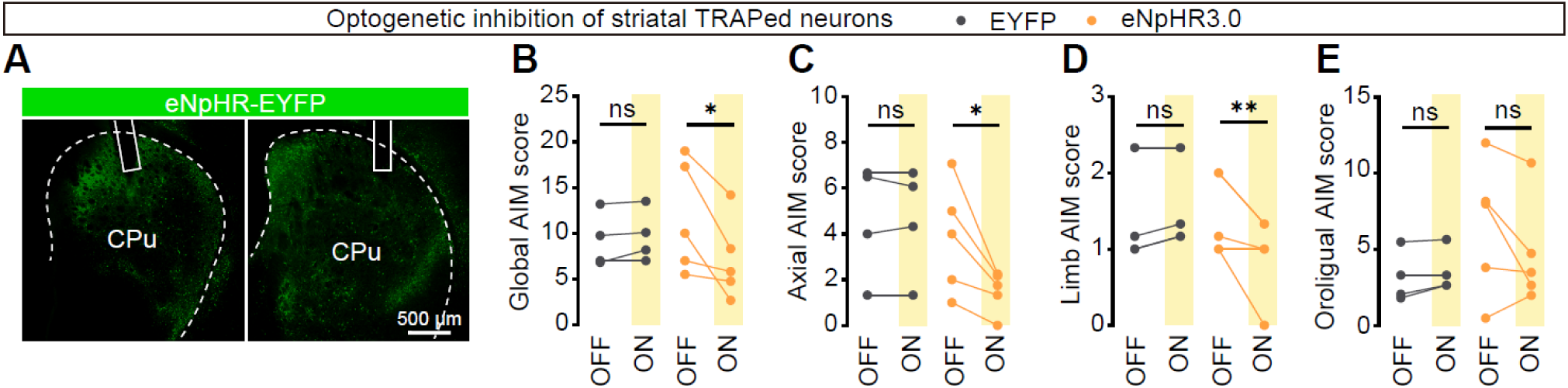
Optogenetic inhibition of dyskinesia-linked striatal ensembles suppresses ongoing LID. (A) Representative coronal sections showing Cre-dependent eNpHR3.0–EYFP expression in the striatum of TRAP2 mice (TRAPed ensemble neurons) and the corresponding optical fiber placements above two CPu sites. (B–E) AIM score quantified during interleaved light-OFF and light-ON epochs in EYFP controls (gray) and eNpHR3.0 mice (orange). (B) Global, F_(1, 7)_ = 7.470, \**p* = 0.0292; EYFP, *p* = 0.9277; eNpHR3.0, \**p* = 0.0153. (C) Axial, F_(1, 7)_ = 7.233, \**p* = 0.0311; EYFP, *p* = 0.9993; eNpHR3.0, \*\**p* = 0.0094. (D) Limb, F_(1, 7)_ = 8.894, \**p* = 0.0204; EYFP, *p* = 0.6989; eNpHR3.0, \**p* = 0.0178. (E) Orolingual, F_(1, 7)_ = 2.555, *p* = 0.154; EYFP, *p* = 0.9131; eNpHR3.0, *p* = 0.1741. EYFP, *N* = 4; eNpHR3.0, *N* = 5. Two-way ANOVA followed by Sidak’s post hoc test. *N*, animals. Data are mean ± SEM.

**Fig. S4:**
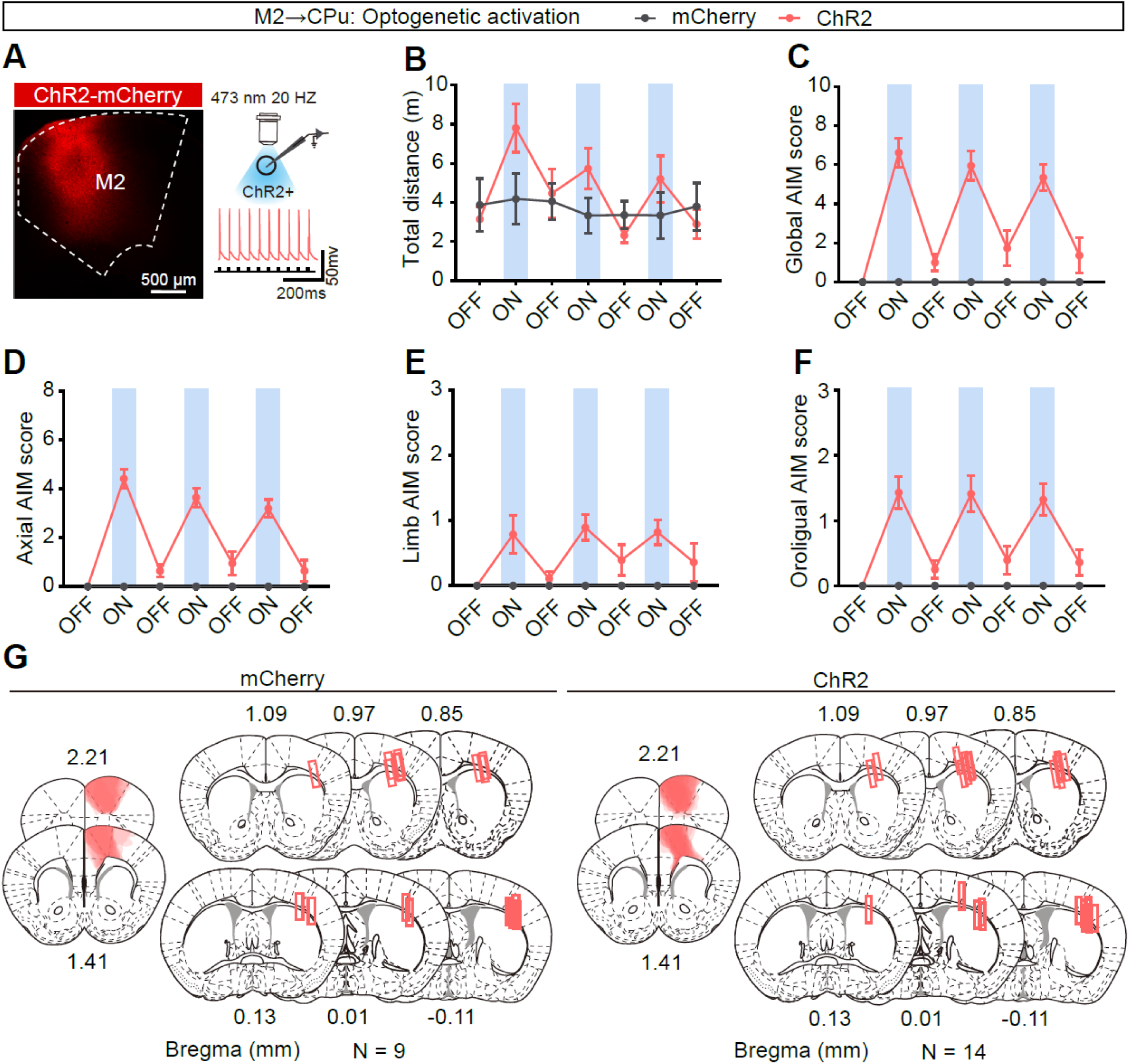
Projection-wide M2 terminal stimulation induces dyskinesia-like behavior in the absence of levodopa. (A) Left, representative image showing ChR2-mCherry expression in M2. Right, *ex vivo* validation of functional opsin expression: whole-cell recording in M2 showing light-evoked spiking in a ChR2-expressing M2 neuron. (B–F) Trial-resolved behavioral dynamics underlying the averaged OFF–ON comparisons shown in Fig. 1. Plots show distance traveled (B) and AIM score (c–f; global, axial, limb, and orolingual) across three consecutive OFF–ON stimulation cycles delivered to M2 axon terminals in striatum in the absence of levodopa. Blue shading denotes light ON epochs. mCherry, *N* = 9; ChR2, *N* = 14. (G) Histological verification of viral expression and implant targeting. Left, overlay of M2 viral expression across animals (shaded). Right, reconstructed optical fiber placements in CPu (rectangles) mapped onto atlas sections. *N* = 14. *N*, animals. Data are mean ± SEM.

**Fig. S5:**
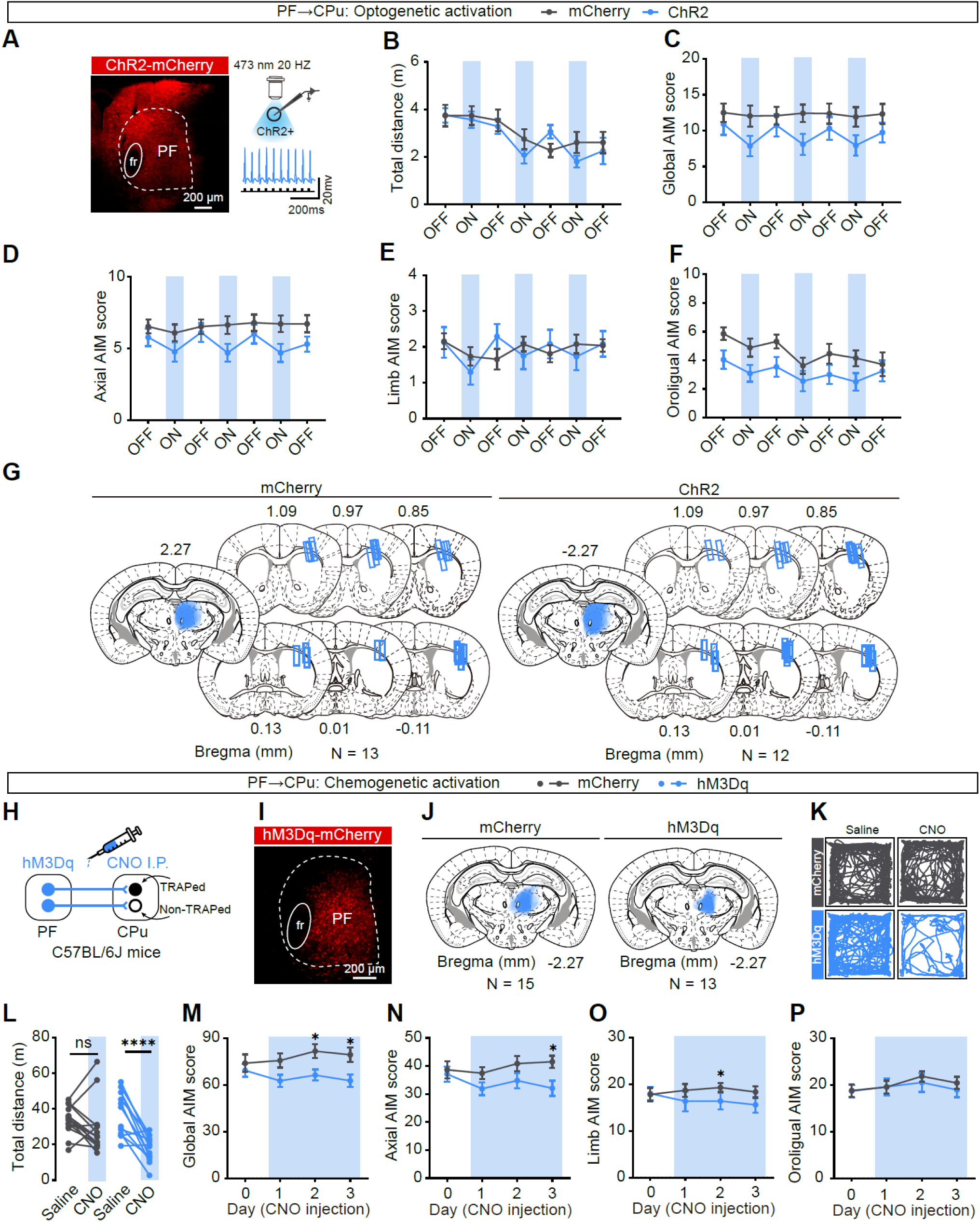
Projection-wide PF activation suppresses dyskinesia through both optogenetic and chemogenetic approaches. (A) Left, representative image showing ChR2-mCherry expression in PF. Right, *ex vivo* validation of functional opsin expression: whole-cell recording in PF showing light-evoked spiking in a ChR2-expressing PF neuron. (B–F) Trial-resolved behavioral dynamics underlying the averaged OFF–ON comparisons shown in Fig. 1. Plots show distance traveled (B) and AIM scores (C–F; global, axial, limb, and orolingual) across three consecutive OFF–ON stimulation cycles delivered to PF axon terminals in striatum in the absence of levodopa. Blue shading denotes light ON epochs. mCherry, *N* = 13; ChR2, *N* = 12. (G) Histological verification of viral expression and implant targeting. Left, overlay of PF viral expression across animals (shaded). Right, reconstructed optical fiber placements in CPu (rectangles) mapped onto atlas sections. *N* = 12. (H) Schematic of chemogenetic activation of PF→CPu pathway by hM3Dq and intraperitoneal clozapine-N-oxide (CNO). (I) Representative hM3Dq-mCherry expression in PF. (J) Summary of hM3Dq expression across animals (blue shading). *N* = 13. (K) Representative open field trajectories after saline or CNO administration (20 min) for mCherry controls and hM3Dq-expressing mice. (L) Total distance traveled following CNO injection in each group. F_(1, 26)_ = 9.511, \*\**p* = 0.0023; mCherry, *N* = 15, *p* > 0.9999; ChR2, *N* = 13, \*\*\*\**p* < 0.0001. Two-way ANOVA followed by Sidak’s post hoc test. (M–P) Global (M), axial (N), limb (O), and orolingual (P) AIM score measured across repeated daily CNO sessions. (M) Global, F_(3, 78)_ = 1.983, *p* = 0.1234; Day 0, *p* = 0.9434; Day 1, *p* = 0.1445; Day 2, \**p* = 0.0477; Day 3, \**p* = 0.049. (N) Axial, F_(3, 78)_ = 2.775, \**p* = 0.0468; Day 0, *p* = 0.9469; Day 1, *p* = 0.1551; Day 2, *p* = 0.1796; Day 3, \**p* = 0.0102. (O) Limb, F_(3, 78)_ = 1.983, *p* = 0.1358; Day 0, *p* = 0.992; Day 1, *p* = 0.1523; Day 2, \**p* = 0.0372; Day 3, *p* = 0.1752. (P) Orolingual, F_(3, 78)_ = 0.7326, *p* = 0.7326; Day 0, *p* = 0.9671; Day 1, *p* = 0.8515; Day 2, \**p* = 0.3638; Day 3, *p* = 0.6435. mCherry, *N* = 15, hM3Dq, *N* = 13. Two-way ANOVA followed by Sidak’s post hoc test. *N*, animals. Data are mean ± SEM.

**Fig. S6:**
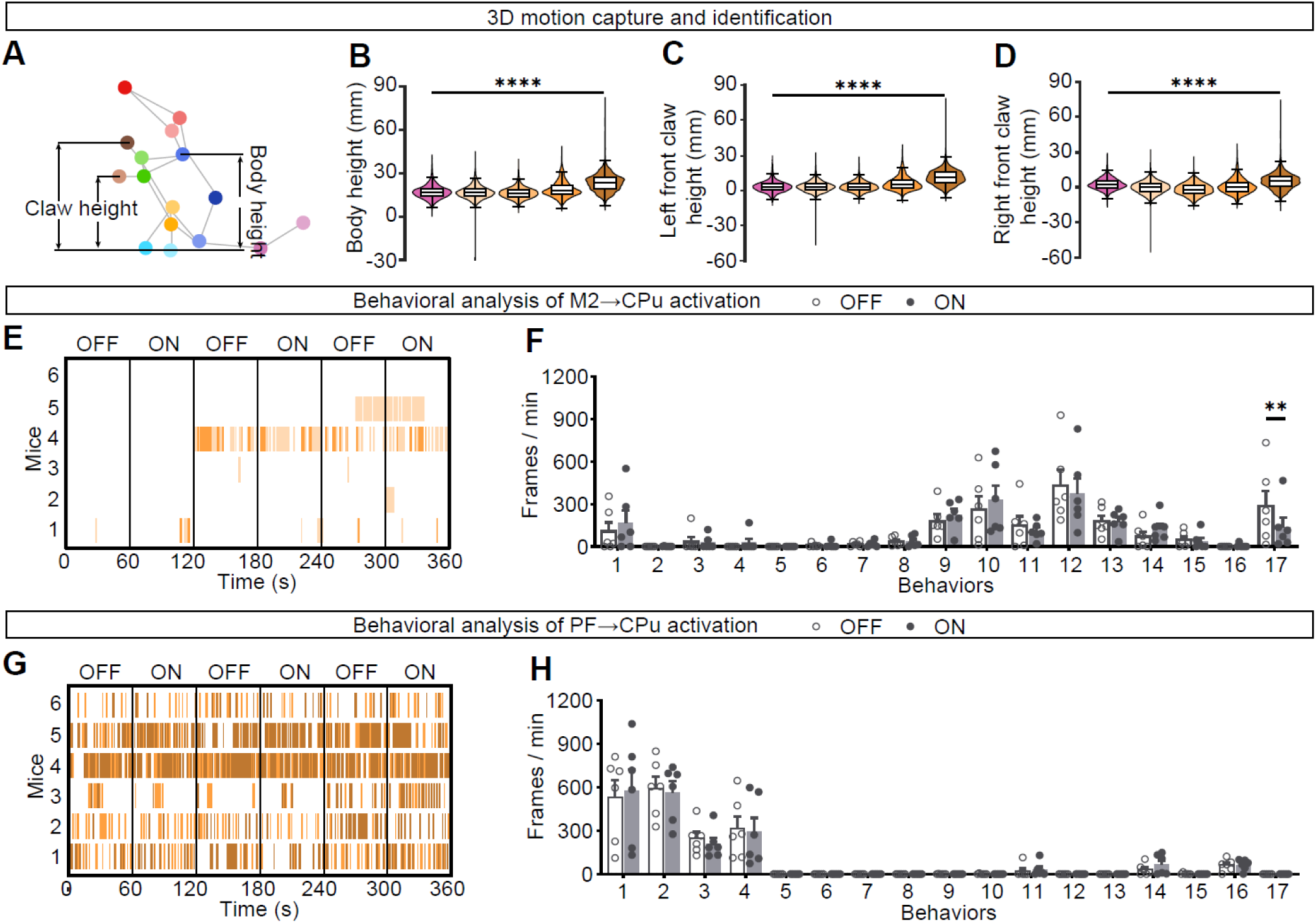
Additional kinematic features validate dyskinesia phenotypes and control analyses in mCherry mice. (A) Definition of vertical kinematic features extracted from the reconstructed 3D skeleton. Body height was defined as the vertical distance from the neck point to the reference plane set by the mean height of the two hind paws. Left and right forepaw heights were defined analogously as the vertical distance from each forepaw to the mean height of the two hind paws. (B–D) Distributions of body height (B), left forepaw height (C), and right forepaw height (D) for Walking and four LID-associated phenotypes (LID_A1–LID_A4, as defined in Figure 2b) pooled from all animals included in the optogenetic cohorts. (B) Body height, F_(4, 138194)_ = 8920.48, \*\*\*\**p* < 0.0001; All groups compared with others, \*\*\*\**p* < 0.0001. (C) Left front claw height, F_(4, 138194)_ = 11354.66, \*\*\*\**p* < 0.0001; Walking vs. LID_A1, *p* = 0.056; Walking vs. LID_A2, *p* = 0.5619; LID_A1 vs. LID_A2, *p* = 0.6615; All others, \*\*\*\**p* < 0.0001. (D) Right front claw height, F_(4, 138194)_ = 5586.84, \*\*\*\**p* < 0.0001; All groups compared with others, \*\*\*\**p* < 0.0001. Walking, *n* = 16,172; LID_A1, *n* = 52,225; LID_A2, *n* = 31,260; LID_A3, *n* = 19,576; LID_A4, *n* = 18,971, *N* = 24. Two-way ANOVA followed by Tukey’s post hoc test. (E, G) Ethograms from mCherry control mice during striatal photostimulation of M2→CPu inputs (E) or PF→CPu inputs (G), plotted for the LID-associated phenotype highlighted in Fig. 2G (E) and Fig. 2I (G), respectively. Colored ticks indicate frames classified as the corresponding LID phenotype across three alternating OFF/ON epochs (60 s each). Each row represents one mouse. (F, H) Occupancy of the 17 BehaviorAtlas phenotypes (IDs as in Fig. 2B), quantified as frames per min during light OFF versus ON epochs in mCherry control mice with stimulation of M2→CPu inputs (F) or PF→CPu inputs (H). OFF and ON values were computed within the same animals across the three cycles shown in (E, H). (F) M2→CPu, F_(16, 85)_ = 1.859, \**p* = 0.0361; Behavior 17, \*\**p* = 0.0024. (H) PF→CPu, F_(16, 85)_ = 1.405, *p* = 0.1592. *N* = 6. Two-way ANOVA followed by Sidak’s post hoc test. *n*, frames; *N*, animals. Data are means ± SEMs or box-and-whisker plots.

**Fig. S7:**
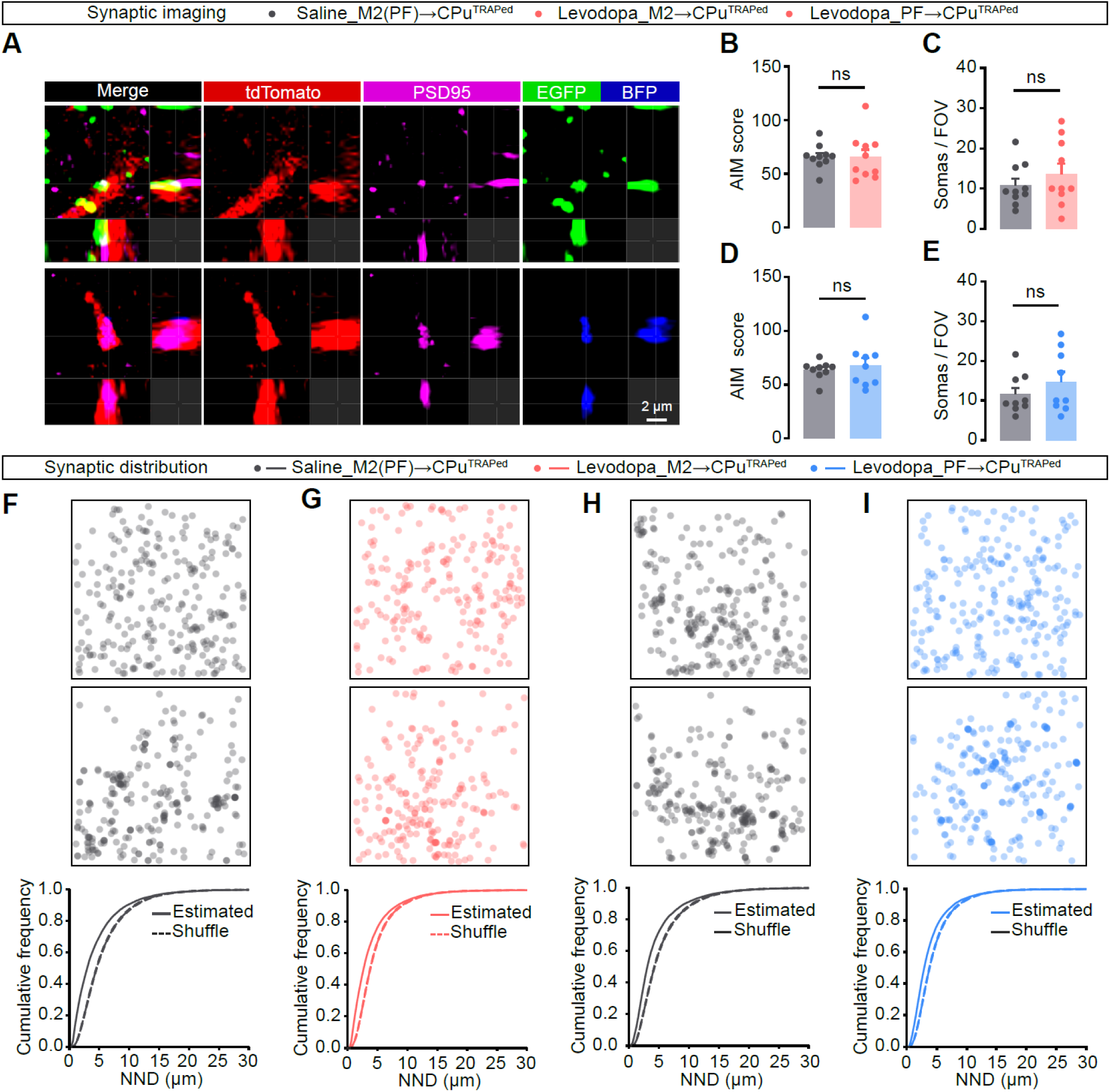
M2 and PF contacts onto dyskinesia-linked ensemble neurons exhibit non-random spatial organization. (A) Representative orthogonal view from confocal z-stacks illustrating the criteria used to define putative synaptic contacts onto FosTRAP-tagged striatal neurons. tdTomato^+^ TRAPed neuron surface (red) and PSD95 puncta (magenta) are shown together with presynaptic boutons labeled by synaptophysin reporters (synaptophysin-EGFP for M2 terminals, green; top; synaptophysin-mTagBFP2 for PF terminals, blue; bottom). (B, C) Cohort validation for M2→CPu^TRAPed^ synapse imaging. Saline, *N* = 10; Levodopa, *N* = 10. (B) Global AIM score did not differ between saline- and levodopa-maintained groups used for M2 synapse imaging. ns, p = 0.9777, two-tailed unpaired Student’s *t* test. (C) The number of tdTomato^+^ somata sampled per field of view (FOV) was comparable across groups. ns, p = 0.3764, two-tailed unpaired Student’s *t* test. (D, E) Same as (B, C) for PF→CPu^TRAPed^ synapse imaging. Saline, *N* = 9; Levodopa, *N* = 9. (D) Global AIM score. ns, *p* = 0.5677; two-tailed unpaired Student’s *t* test. (E) Number of tdTomato^+^ somata. ns, *p* = 0.3315; two-tailed unpaired Student’s *t* test. (F–I) Nearest-neighbor distance (NND) analysis comparing observed synapse locations (“Estimated”) with within-FOV randomized controls (“Shuffle”). For each condition, the top panels show representative maps of estimated synapse coordinates; the middle panels show shuffled maps generated by reassigning synapses to postsynaptic sites (PSD95 puncta on tdTomato^+^ surfaces) within the same FOV (1,000 randomizations; see Methods); the bottom panels show cumulative NND distributions for estimated versus shuffled synapses. The dashed line represents the 95% confidence interval of randomized cumulative distribution. (F) M2→CPu^TRAPed^ synapses, Saline. *n* = 17,356, *N* = 10. \*\*\*\**p* < 0.0001; Kolmogorov-Smirnov test. (G) M2→CPu^TRAPed^ synapses, levodopa-maintained. *n* = 19,005, *N* = 10. \*\*\*\**p* < 0.0001; Kolmogorov-Smirnov test. (H) PF→CPu^TRAPed^ synapses, saline-maintained. *n* = 16,453, *N* = 9. \*\*\*\**p* < 0.0001; Kolmogorov-Smirnov test. (I) PF→CPu^TRAPed^ synapses, levodopa-maintained. *n* = 18,153, *N* = 9. \*\*\*\**p* < 0.0001; Kolmogorov-Smirnov test. *n*, synapses; *N*, animals. Data are mean ± SEM.

**Fig. S8:**
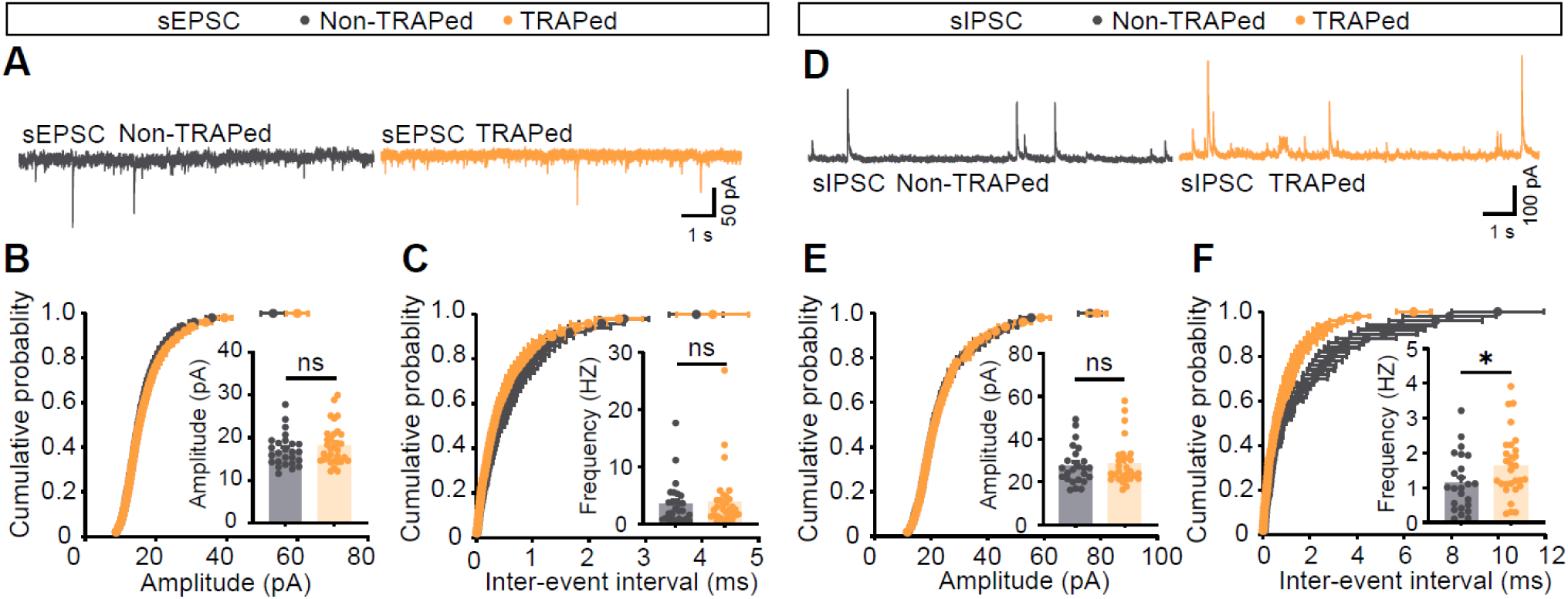
Dyskinesia-linked ensemble neurons show increased spontaneous inhibitory, but not excitatory, synaptic drive. (A) Representative spontaneous excitatory postsynaptic current (sEPSC) traces recorded from neighboring tdTomato^−^ non-TRAPed (gray) and tdTomato^+^ TRAPed (orange) neurons in the dorsolateral CPu of LID mice. (B, C) Cumulative distributions of sEPSC amplitudes (B) and inter-event intervals (C); insets summarize per-cell mean amplitude and event frequency. (B) Amplitude, ns, *p* = 0.3311. (C) Frequency, ns, *p* = 0.7593. NonTRAPed, *n* = 26, *N* = 9; TRAPed, n = 32, *N* = 9. Mann-Whitney U test. (D) Representative spontaneous inhibitory postsynaptic current (sIPSC) traces from non-TRAPed and TRAPed neurons. (E, F) Cumulative distributions of sIPSC amplitudes (E) and inter-event intervals (F); insets summarize per-cell mean amplitude and event frequency. (E) Amplitude, ns, *p* = 0.7465, Mann-Whitney U test. (F) Frequency, \**p* = 0.0466, two-tailed unpaired Student’s *t* test. NonTRAPed, *n* = 24, *N* = 9; TRAPed, *n* = 29, *N* = 9. *n*, cells; *N*, animals. Data are mean ± SEM.

**Fig. S9:**
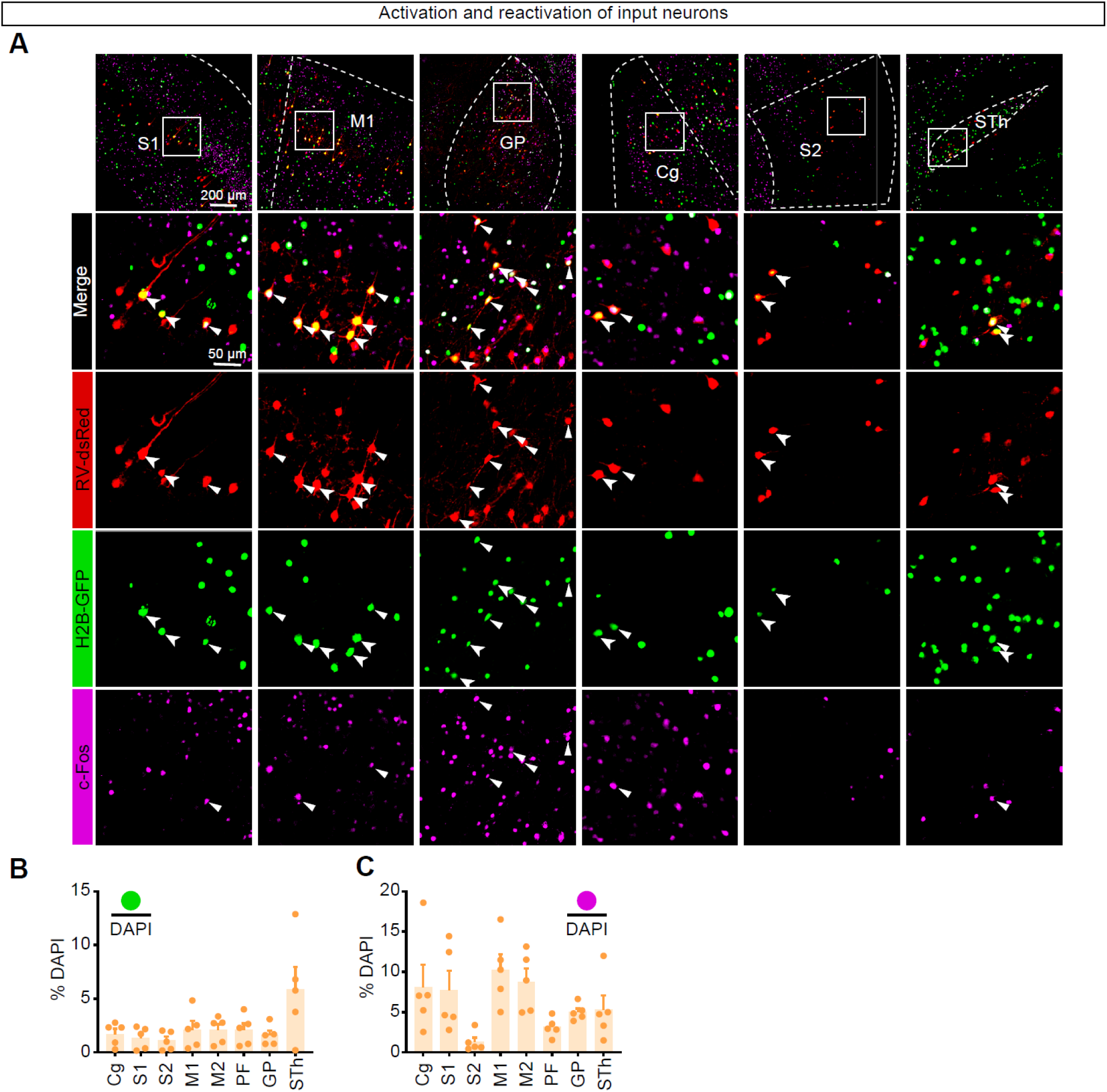
Activity states of rabies-defined input neurons across additional upstream regions during dyskinesia. (A) Representative coronal sections showing monosynaptic inputs (RV-dsRed, red) to striatal TRAPed starter neurons and their activity state across two LID episodes in representative cortical and basal ganglia regions (S1, M1, S2, Cg, GP, and STh). TRAP-captured activation during the first LID episode is marked by nuclear TRAP-H2B-GFP (green), and reactivation during the second LID episode is marked by c-Fos immunoreactivity (magenta). Top row, low-magnification views with regions of interest indicated; bottom rows, higher-magnification views of the boxed areas. Arrowheads denote activated input neurons (RV-dsRed^+^H2B-GFP^+^); triangles denote reactivated input neurons (RV-dsRed^+^H2B-GFP^+^c-Fos^+^). (B, C) Density of TRAP-H2B-GFP^+^ nuclei (B, *N* = 5) and c-Fos^+^ nuclei (C, *N* = 5) in the individual regions, expressed as the percentage of total DAPI^+^ nuclei (same animals and sampling strategy as in Fig. 5).

**Fig. S10:**
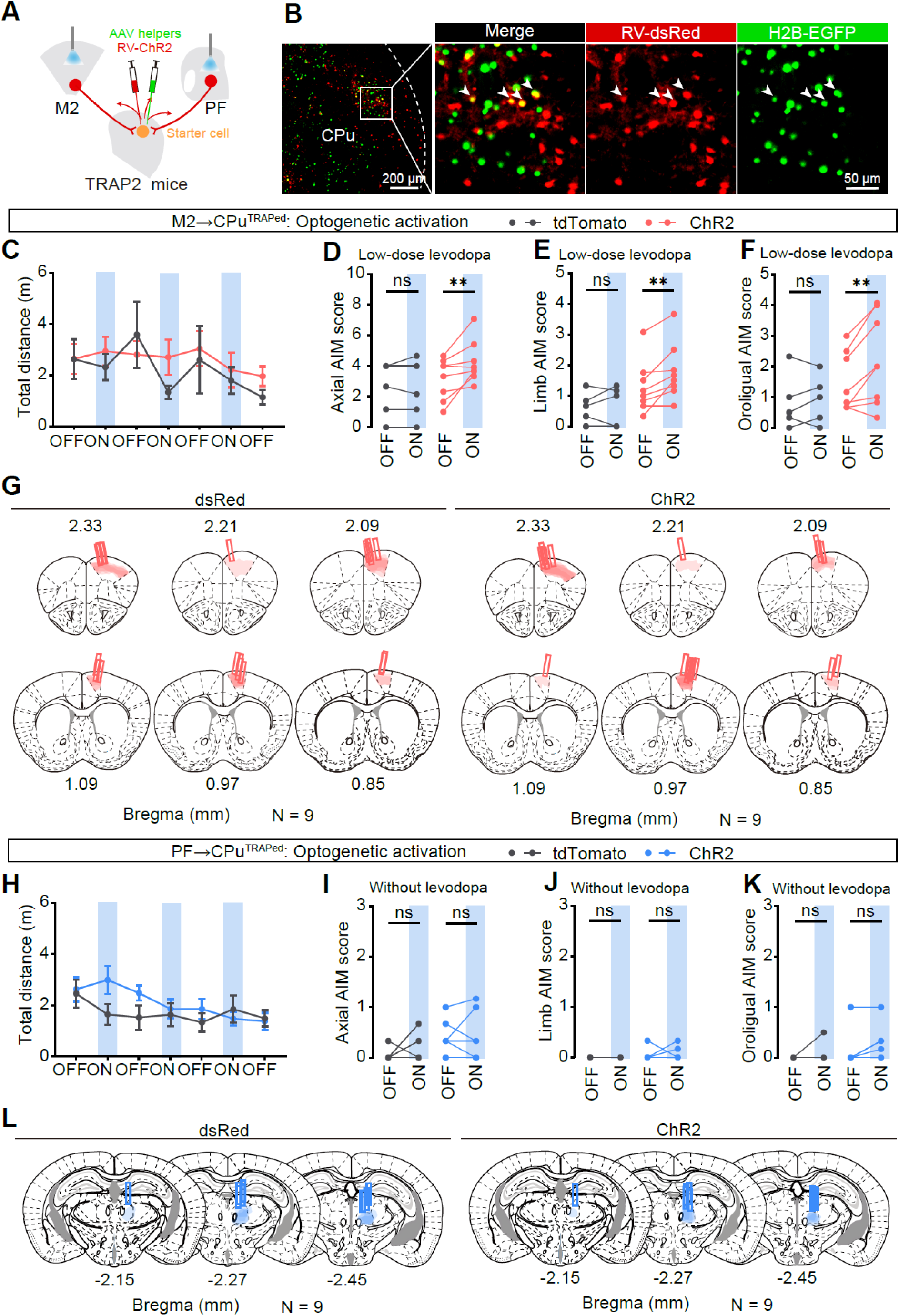
Trial-resolved effects of circuit-restricted stimulation of M2→ensemble and PF→ensemble inputs. (A) Experimental strategy for circuit-restricted optogenetic activation of monosynaptic inputs to striatal TRAPed neurons. In TRAP2 mice, AAV helper viruses were delivered to CPu to enable EnvA-pseudotyped, glycoprotein-deleted rabies labeling of presynaptic input neurons; rabies encoding tdTomato (control) or ChR2-dsRed (ChR2) was used to permit optical stimulation of input populations (M2 or PF) that innervate CPu^TRAPed^ starter cells. (B) Representative images of CPu showing starter cells (arrowheads) identified by co-localization of RV-dsRed (red) and TRAP-dependent H2B-EGFP (green). Left, low magnification; right, enlarged view of boxed area. (C, H) Total distance traveled across successive OFF/ON epochs during optical stimulation of M2→CPu^TRAPed^ (C) or PF→CPu^TRAPed^ (H) inputs. dsRed, *N* = 9; ChR2, *N* = 9. (D–F, I–K) AIM subtype scores during M2→CPu^TRAPed^ (D–F, tdTomato, *N* = 5; ChR2, *N* = 8) or PF→CPu^TRAPed^ (I–K, tdTomato, *N* = 9; ChR2, *N* = 9) stimulation. Axial (D, I), limb (E, J), and orolingual (F, L) AIM scores were quantified during paired OFF and ON epochs for dsRed controls and ChR2 mice. (D, I) Axial AIM score. (D) M2→CPu^TRAPed^, F_(1, 11)_ = 4.552, *p* = 0.0562; tdTomato, *p* = 0.9956; ChR2, \*\**p* = 0.0091. (I) PF→CPu^TRAPed^, F_(1, 16)_ = 1.239, *p* = 0.282; tdTomato, *p* = 0.1878; ChR2, *p* = 0.9806. (E, J) Limb AIM score. (E) M2→CPu^TRAPed^. F_(1, 11)_ = 3.832, *p* = 0.0761; tdTomato, *p* = 0.9149; ChR2, \*\**p* = 0.0077. (J) PF→CPu^TRAPed^. F_(1, 16)_ = 0.1055, *p* = 0.7495; tdTomato, *p* = 0.5651; ChR2, *p* = 0.2816. (F, K) Orolingual AIM score. (F) M2→CPu^TRAPed^. F_(1, 11)_ = 3.667, *p* = 0.0818; tdTomato, *p* = 0.9159; ChR2, \*\**p* = 0.0088. (K) PF→CPu^TRAPed^. F_(1, 16)_ = 0.2467, *p* = 0.6262; tdTomato, *p* = 0.5142; ChR2, *p* = 0.1834. Two-way ANOVA followed by Sidak’s post hoc test. (G, L) Reconstruction of rabies-labeled neurons (shading) and optical fiber placements (rectangles) in M2 (G, *N* = 9) or PF (L, *N* = 9) for animals contributing to (C–F) or (I–K). *N*, animals. Data are mean ± SEM.

**Fig. S11:**
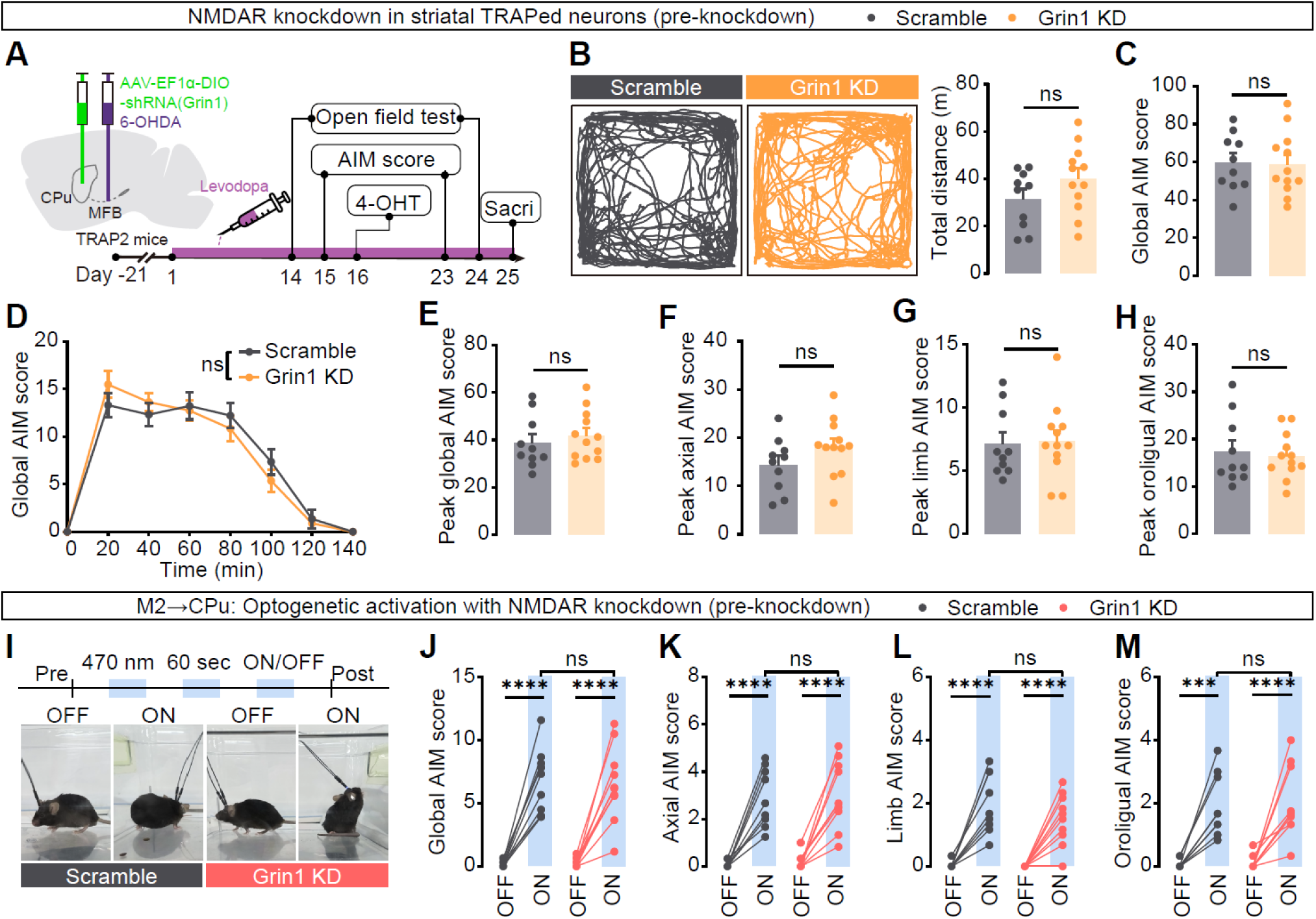
Scramble and *Grin1* knockdown cohorts are matched before activity-dependent knockdown induction. (A) Experimental design and timeline for Cre-dependent delivery of *Grin1* shRNA (or scramble control) to the CPu of TRAP2 mice followed by unilateral 6-OHDA lesioning and levodopa priming. “Pre-knockdown” measurements were collected before 4-hydroxytamoxifen (4-OHT)–triggered TRAP recombination, when Cre-dependent shRNA is not yet expressed in the LID-tagged ensemble. (B) Open-field locomotion measured before 4-OHT induction. Left, representative center-point trajectories (20-min session). Right, total distance traveled. Scramble, *N* = 10; *Grin1* KD, *N* = 12. *p* = 0.1374; two-tailed unpaired Student’s *t* test. (C) Global AIM score during LID before 4-OHT induction. Scramble, *N* = 10; *Grin1* KD, *N* = 12. ns, *p* = 0.9013; two-tailed unpaired Student’s *t* test. (D) Time course of global AIM score following a single levodopa injection before 4-OHT induction. F_(7, 140)_ = 1.113, *p* = 0.3586. Scramble, *N* = 10; *Grin1* KD, *N* = 12. Two-way ANOVA followed by Sidak’s post hoc test. (E–H) Peak AIM score extracted from the 20 ∼ 60 min LID window after levodopa administration, including global (E) and subtype scores for axial (F), limb (G), and orolingual (H) components. (E) Global, *p* = 0.526. (F) Axial, *p* = 0.1517. (G) Limb, *p* = 0.879. (H) Orolingual, *p* = 0.7397. Scramble, *N* = 10; *Grin1* KD, *N* = 12. Two-tailed unpaired Student’s *t* test. (I) Schematic of the optogenetic assay used to probe M2→CPu drive prior to knockdown induction, with representative snapshots during OFF and ON epochs. (J–M) AIM score during OFF versus ON epochs of M2→CPu stimulation before 4-OHT induction, including global (J) and axial (K), limb (L), and orolingual (M) subtypes, demonstrating comparable stimulation-evoked dyskinesia in both cohorts prior to knockdown. (J) Global, F_(1, 17)_ = 0.1113, *p* = 0.7427; Scramble, \*\*\*\**p* < 0.0001; *Grin1* KD, \*\*\*\**p* < 0.0001; Scramble_ON vs. *Grin1* KD_ON, *p* = 0.9353. Two-way ANOVA followed by Sidak’s post hoc test. (K) Axial, F_(1, 17)_ = 0.001, *p* = 0.9746; Scramble, \*\*\*\**p* < 0.0001; *Grin1* KD, \*\*\*\**p* < 0.0001; Scramble_ON vs. *Grin1* KD_ON, *p* = 0.9644. (L) Limb, F_(1, 17)_ = 1.328, *p* = 0.2652; Scramble, \*\*\*\**p* < 0.0001; *Grin1* KD, \*\*\*\**p* < 0.0001; Scramble_ON vs. *Grin1* KD_ON, *p* = 0.1722. (M) Orolingual, F_(1, 17)_ = 0.01671, *p* = 0.8987; Scramble, \*\*\**p* = 0.0002; *Grin1* KD, \*\*\*\**p* < 0.0001; Scramble_ON vs. *Grin1* KD_ON, *p* = 0.9592. Scramble, *N* = 9; *Grin1* KD, *N* = 10. Two-way ANOVA followed by Sidak’s post hoc test. *N*, animals. Data are mean ± SEM.

